# Chemogenetic evidence that rapid neuronal *de novo* protein synthesis is required for consolidation of long-term memory

**DOI:** 10.1101/704965

**Authors:** Prerana Shrestha, Pinar Ayata, Pedro Herrero-Vidal, Francesco Longo, Alexandra Gastone, Joseph E. Ledoux, Nathaniel Heintz, Eric Klann

**Author notes:** These authors contributed equally (Prerana Shrestha, Pinar Ayata). Corresponding senior authors.

## Abstract

Translational control of memory processes is a tightly regulated process where the coordinated interaction and modulation of translation factors provides a permissive environment for protein synthesis during memory formation. Existing methods used to block translation lack the spatiotemporal precision to investigate cell-specific contributions to consolidation of long-term memories. Here, we have developed a novel chemogenetic mouse resource for <u>ce</u>ll type-specific and drug-<u>i</u>nducible <u>p</u>rotein <u>s</u>ynthesis inhibition (ciPSI) that utilizes an engineered version of the catalytic kinase domain of dsRNA-activated protein (PKR). ciPSI allows rapid and reversible phosphorylation of eIF2α causing a block on general translation by 50% *in vivo*. Using this resource, we discovered that temporally structured pan-neuronal protein synthesis is required for consolidation of long-term auditory threat memory. Targeted protein synthesis inhibition in CamK2α expressing glutamatergic neurons in lateral amygdala (LA) impaired long-term memory, which was recovered with artificial chemogenetic reactivation at the cost of stimulus generalization. Conversely, genetically reducing phosphorylation of eIF2α in CamK2α positive neurons in LA enhanced memory strength, but was accompanied with reduced memory fidelity and behavior inflexibility. Our findings provide evidence for a finely tuned translation program during consolidation of long-term threat memories.

## Introduction

Memory is the capacity of an organism to encode, store, and retrieve information, and often guides the action of the organism towards a better survival outcome. Aversive life-threatening events often lead to long-term associative memories between the environment in which those events were experienced in and the threat, such that a salient cue from the event when presented again can elicit species-specific defensive behaviors. Pavlovian cued threat conditioning is a useful experimental paradigm for understanding the biological substrates of an associative threat memory, in which a neutral cue (e.g. light or tone) co-presented with an innately aversive stimulus (e.g. air puff or footshock) elicits a defensive response such as freezing by a repeat presentation by itself^1^. Cued threat conditioning is particularly amenable to studying memory consolidation process because one-trial training is sufficient to form a persistent long-term memory and a unimodal cue can be used for memory retrieval. Long-term aversive memories are thought to recruit a distributed network of neurons across the brain, including subnuclei within amygdala, hippocampus, thalamus, as well as the neocortex, depending on the brain state, cue complexity, and the sensory pathways engaged^2, 3^. Lateral amygdala is the central sensory gateway for the amygdaloid complex and is crucially engaged both in the processing and storage of the associative aversive memories^4, 5, 6^.

Decades of studies have reported that consolidation of long-term memories requires one or more waves of new protein synthesis after the memory is first encoded^7, 8, 9, 10^; however, the use of pharmacological protein synthesis inhibitors (PSI) in these studies is burdened with limitations. For instance, an extensively used PSI, anisomycin, causes unwanted side effects that include superinduction of immediate early genes, activation of stress signaling pathways, and catecholamine release^11, 12^. Pharmacological PSIs also do not discriminate between molecularly heterogeneous cell populations and thus, do not allow manipulation of cell autonomous protein synthesis. Genetic approaches to alter the translation machinery at the initiation step provide less ambiguous evidence for the role of protein synthesis in memory processes^13, 14^. However constitutive deletion of genes encoding translational control molecules lacks the temporal control needed to convincingly interrogate the role of protein synthesis in memory consolidation. Past attempts at making chemogenetic tools for blocking protein synthesis have had limited success^15^.

A key regulatory step in mammalian protein synthesis is the assembly of active eIF2.GTP and initiator-methionyl tRNA into a ternary complex that binds the 40S ribosome. Various types of cellular stress engage pathway-specific protein kinases that phosphorylate the α subunit of eIF2 on serine 51, which converts eIF2 from a substrate to a competitive inhibitor of eIF2B, a guanine exchange factor that promotes the conversion of inactive eIF2.GDP to active eIF2.GTP. Thus, phosphorylation of eIF2α stops the recycling of the ternary complex and inhibits general translation^16^. However, phosphorylation of eIF2α also leads to a paradoxical increase in gene-specific translation of transcripts harboring upstream open reading frames (uORFs) by overcoming the inhibitory effects of uORFs on reinitiation at the putative start codon as ternary complexes decline^17^. Dephosphorylation of eIF2α occurs during L-LTP as well as during consolidation and reconsolidation of associative memories, and is thought to remove the constraint on general translation^14, 18^. Here, we have developed a novel cell type-specific drug-inducible protein synthesis inhibitor (ciPSI) that utilizes an inducible form of the kinase domain of double-stranded RNA activated protein kinase (iPKR) as an actuator for phosphorylating endogenous eukaryotic initiation factor 2α on serine 51 (S51). This resulted in 50% reduction in *de novo* translation followed by rapid clearance of iPKR and p-eIF2α within hours after induction in awake behaving mice. We investigated the time-limited role of protein synthesis in pan-neuronal Nestin.iPKR mice and subsequently in Ca^2+^/calmodulin-dependent protein kinase 2α (CamK2α+)-expressing glutamatergic neurons in the lateral amygdala (LA) during consolidation of auditory threat memory. We found that long-term threat memories are particularly labile to pan-neuronal and LA CamK2α+ cell type-specific protein synthesis disruption in the first hour after training. We further found that the translation program regulated by phosphorylation status of eIF2α plays an important role in calibrating memory strength and fidelity. Our findings provide new mechanistic insight into the nature of long-term threat memory consolidation.

## Results

### A chemogenetic resource for cell type-specific protein synthesis inhibition

To induce phosphorylation of eIF2α, we engineered the kinase domain of the eIF2α kinase protein kinase RNA-activated (PKR), to harbor a recognition site (NS5A/B) for a recombinant non-structural protein 3/4 (NS3/4) protease, which can be chemically inhibited with Asunaprevir (ASV)^19, 20^ (Fig. 1a). We first tested toxicity of ASV in amygdala slices from wild-type mice infused with increasing dose of ASV (0, 10 nM, 100 nM and 1 μM) and determined that for all doses tested, there was no significant induction of either the immediate early gene (IEG) cFos or the integrated stress response as assessed by examination of the level of phosphorylated eIF2α (Supplemental Fig. 1a). The kinase domain of PKR is a caspase-generated fragment that is constitutively active and does not require either double-stranded RNA or dimerization for activation^21^. We designed our ciPSI multicistronic construct with NS3/4 protease^22^, EGFP, and inducible PKR (iPKR) transgenes separated by self-cleaving 2A proteinase that allows translation of individual elements by ribosome skipping^23^ (Supplemental Fig. 1b). To test the inhibition of global translation *in vitro*, newly synthesized proteins were metabolically labeled with S^35^ methionine after starvation. In cells transfected with iPKR and NS3/4A, iPKR was degraded by NS3/4 and *de novo* translation was equivalent to the control cells, whereas cells transfected with iPKR plasmid alone in the absence of NS3/4A had substantially reduced translation by about 60%. Unmodified PKR kinase domain (PKRk) similarly reduced *de novo* translation relative to controls, but the PKRk levels were non-responsive to NS3/4 protease (Supplementary Fig. 1c). As predicted, PKRk fragment was detected only in lysates from cells transfected with either unmodified PKRk or modified iPKR (Supplemental Fig. 1d).

We next knocked in the ciPSI multicistronic cassette into the first intron of the mouse *Eef1a1* genomic locus^24^ with two modifications (Fig. 1b and Supplementary Fig. 2a and b). First, EGFP was substituted with an EGFP-L10 fusion to use as a fluorescent marker of cells expressing ciPSI and to eventually enable translating ribosome affinity purification (TRAP) profiling^25^. Second, a STOP cassette flanked by *loxP* sites preceded the ciPSI cassette to allow cell type-specific disruption of protein synthesis when combined with Cre driver mouse lines and/or Cre-expressing viruses. These modifications had no effect on expression of iPKR (data not shown). A mouse Nestin (*Nes*) Cre-driver line^26^ was bred with the ciPSI line to generate transheterozygote Nes.iPKR pan-neuronal ciPSI knock-in mice, which were viable and fertile. The Nes-iPKR mice expressed EGFP-L10 in all neurons in the amygdala, as marked by complete overlap with NeuN staining (Fig. 1d), as well as in cortical areas such as the anterior cingulate cortex and somatosensory cortex, and hippocampal areas, CA1 and CA3, and the dentate gyrus (Supplemental Fig. 2c-g). Despite heterozygous deletion of *Eef1a1* allele and expression of the multicistronic cassette in Nes.iPKR mice, the mice displayed normal spontaneous locomotion as well as thigmotaxis in open field tests (Supplemental Fig. 3a-c).

**Figure 1.**
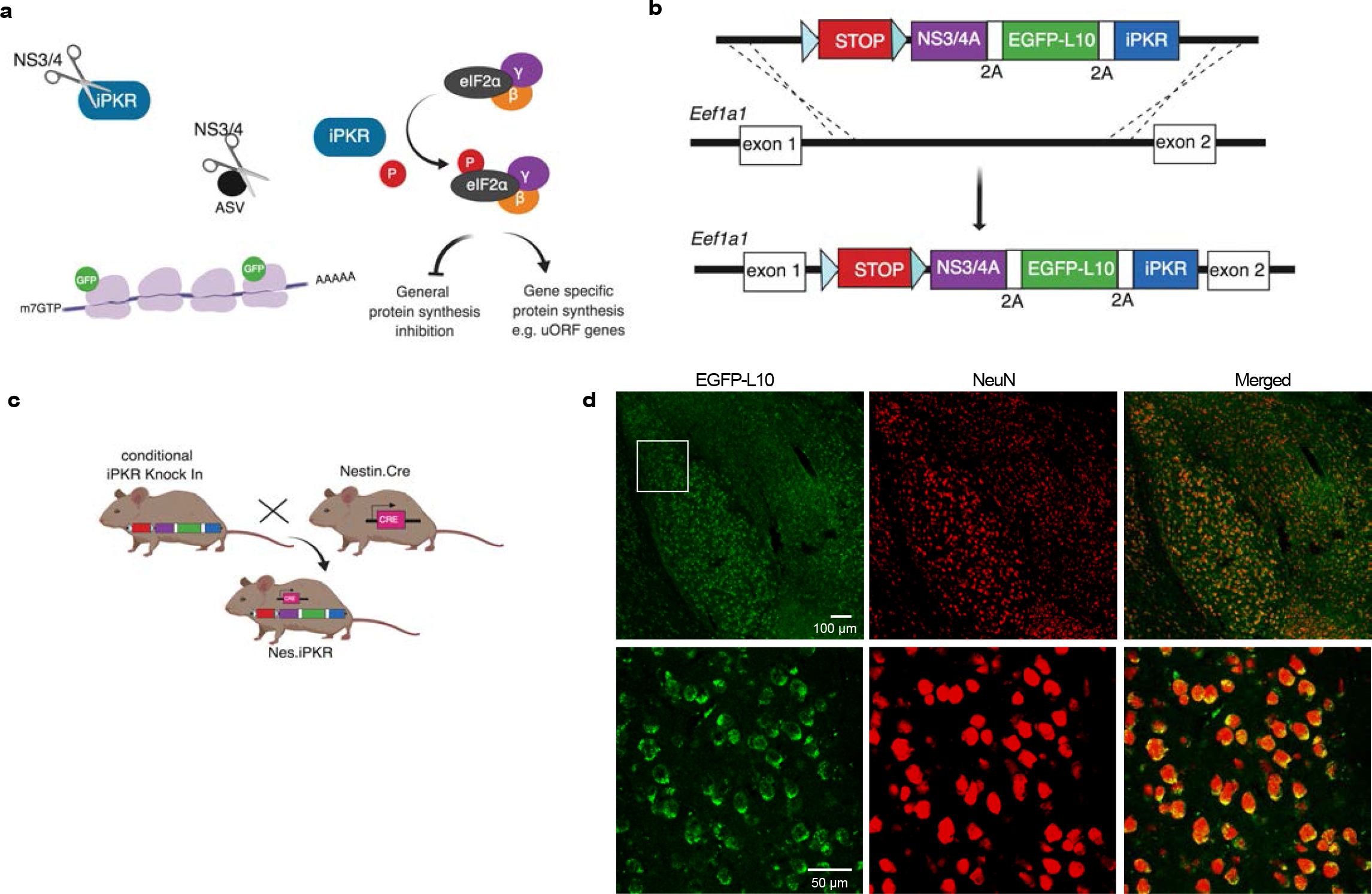
Generation of a chemogenetic resource for cell type-specific protein synthesis inhibition. a) Diagram of the mechanism of action of ciPSI. NS3/4 protease degrades the iPKR kinase domain by binding at the engineered NS5A-5B binding site. NS3/4 inhibitor drug, Asunaprevir (ASV), blocks the activity of NS3/4 protease, thereby releasing the protein synthesis inhibitor iPKR that phosphorylates eIF2α at S51, resulting in an inhibition of general translation while permitting gene-specific translation of transcripts harboring an upstream open reading frame (uORF). b) Schematic of the knock-in transgenesis approach for the ciPSI iPKR cassette into the *Eef1a1* genomic locus. c) Pan neuronal Nes.iPKR mice were generated by breeding Nes.Cre and iPKR knock in mice. d) GFP-tagged ribosomal protein L10 is present in the soma of all neurons in the amygdala, and completely overlaps with NeuN expression. Representative immunofluorescence image shows expression of GFP-L10a (green) in NeuN+ (red) neurons. Inset in the first row is magnified in second row. Scale 100 μm (top), or 50 μm (bottom).

To test the efficiency of ciPSI in blocking protein synthesis *ex vivo*, amygdala slices of Nes.iPKR mice and wild-type (WT) mice were subjected to bio-orthogonal non-canonical amino-acid tagging (BONCAT)^27^ of newly synthesized proteins. Nes.iPKR amygdala slices treated with 1 μM ASV exhibited a sharp decline in *de novo* translation (~20%) compared with controls (Fig. 2a). Because BONCAT uses azidohomoalanine (AHA), a synthetic methionine analog, which can get outcompeted by endogenous methionine *in vivo*, this method does not sufficiently label *de novo* translation *in vivo*. Thus, an independent method of labeling *de novo* translation, surface sensing of translation (SUnSET)^28^, that measures translation elongation and is amenable for *in vivo* labeling, was used to test ciPSI efficiency in awake behaving mice. SUnSET immunoblot showed that Nes.iPKR mice centrally infused with 150 pg ASV exhibited a robust decrease in protein synthesis (~50%) compared to controls (Fig. 2b). The inhibition of protein synthesis was concomitant with a specific increase in phosphorylation of eIF2α at 1h post ASV treatment both *ex vivo* (Fig. 2c) and *in vivo* (Fig. 2d), with no effect on either the extracellular signal regulated kinase (ERK1/2) or mechanistic target of rapamycin (mTORC1) pathways (Fig. 2d). Next, we investigated the ASV pharmacokinetics in Nes.iPKR amygdala lysates by harvesting tissue at increasing time points after drug infusion. We found that peak expression of iPKR is reached at 0.5h, which steadily declines at 1h and 3h and is completely degraded by 6h (Fig. 2e). Subsequently, phosphorylation of eIF2α steadily increased at 0.5h and 1h and then declined to baseline at 3h. As a result of phosphorylation of eIF2α, proteins whose transcripts harbor uORFs, specifically ATF4 and GADD34, accumulated and remained at higher levels compared to control until 3h (Fig. 2e). GADD34 is the regulatory subunit of the eIF2α phosphatase^17^, and the negative feedback due to increased GADD34 levels combined with the degradation of iPKR by NS3/4 protease at 3h time point might be responsible for reduced phosphorylation of eIF2α below the control (Fig. 2e). We also probed for cFos to assess whether cFos levels change with ciPSI, and found that cFos levels significantly decrease below baseline at 3h and 6h following ASV treatment (Fig. 2e) indicating general translation suppression.

**Figure 2.**
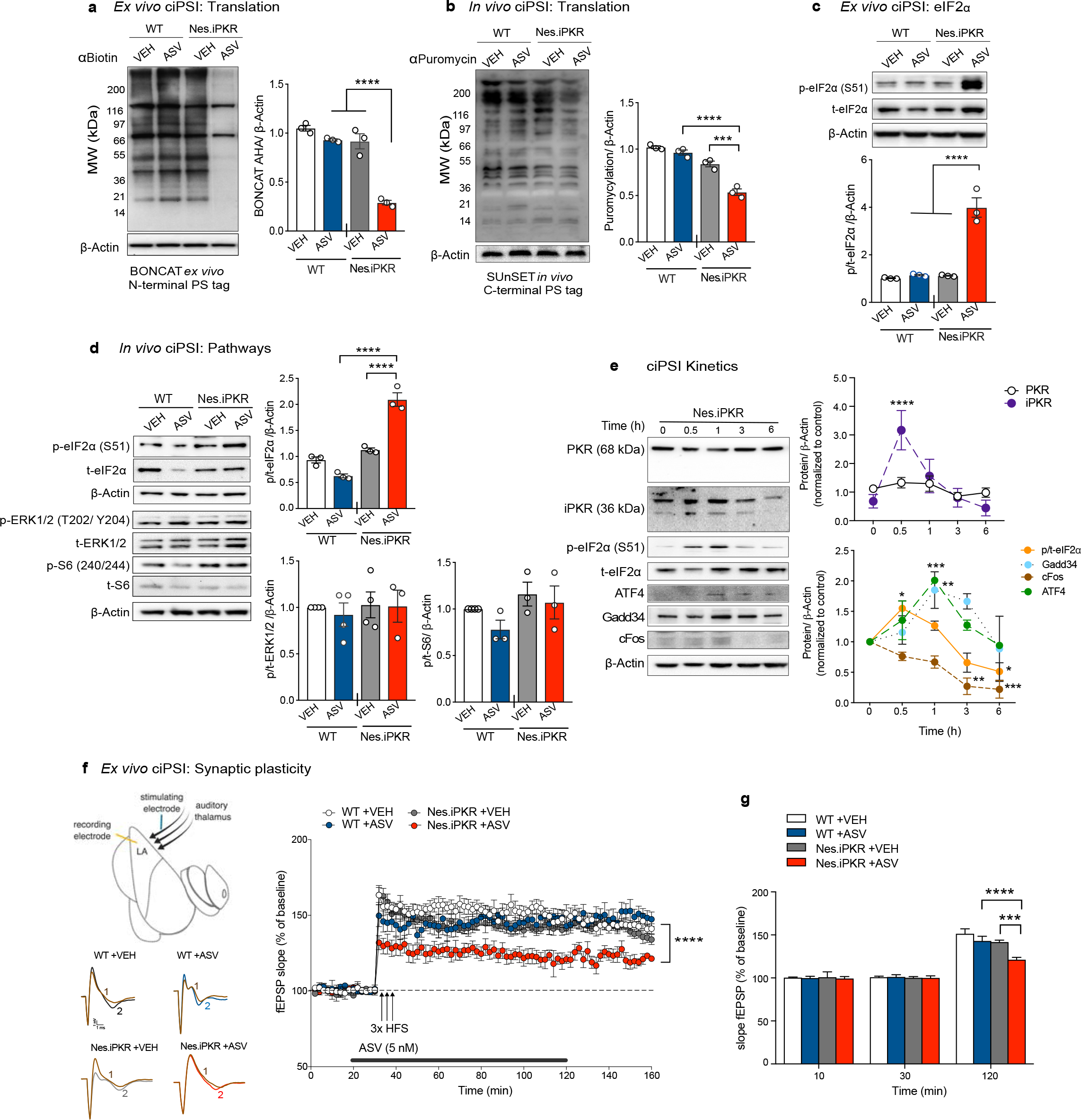
Drug-induced neuronal protein synthesis inhibition. a) *De novo* translation labeled at the N-terminus using BONCAT in Nes.iPKR amygdala slices showed a robust decrease in translation in mutant amygdala treated with 1 μM Asunaprevir (ASV) compared to vehicle (VEH)-treated mutant amygdala (****p<0.0001) as well as ASV-treated wild-type amygdala (****p<0.0001). n = 3 biological replicate lysates, 3 mice per group. Two-way ANOVA with Bonferroni’s post-hoc test. Genotype X Drug interaction: F(1,8) = 34.6, ***p=0.0004; Drug: F(1,8) = 75.2, ****p<0.0001; Genotype: F(1,8) = 80.7, ****p<0.0001. b) *De novo* translation labeled at the C-terminus in awake behaving mice using SUnSET also showed a significant decrease in translation in mutant mice infused with 150 pg ASV (100 nM in 2 μl) compared to vehicle (VEH) infusion (***p<0.001) as well as wild-type (WT) mice infused with ASV (****p<0.0001). n=3 biological replicate lysates, 3 mice per group. Two-way ANOVA with Bonferroni’s post-hoc test. Genotype X Drug interaction: F(1,8) = 19.49, **p=0.0022; Drug: F(1,8) = 41.51, ***p=0.0002; Genotype: F(1,8) = 117.9; p<0.0001. c) Bath application of 1 μM ASV caused a robust phosphorylation of eIF2α in mutant Nes.iPKR amygdala slices compared to ASV-treated wild-type slices and VEH-treated Nes.iPKR slices (****p<0.0001).). n=3 biological replicate lysates, 3 mice per group. Two-way ANOVA with Bonferroni’s post-hoc test. Genotype X Drug interaction: F(1,8) = 44.7, ***p=0.0002; Drug: F(1,8) = 53.5, ****p<0.0001; Genotype: F(1,8) = 51.7, ****p<0.0001. d) ASV infusion *in vivo* (150 pg or 100 nM in 2 μl) also caused a significant increase in p-eIF2α in Nes.iPKR amygdala compared to controls (****p<0.0001). n=3 biological replicate lysates, 3 mice per group. Two-way ANOVA with Bonferroni’s post-hoc test. Genotype X Drug ineraction: F(1,8) = 70.87, ****p<0.0001; Drug: F(1,8) = 19.04, **p<0.0024; Genotype: F(1,8) = 120.4, ****p<0.0001. Major intracellular signaling pathways, ERK1/2 MAPK and mTORC1, assessed by examining p-ERK1/2 (T202/Y204) and p-S6 (S240/244) levels, were unchanged by ASV treatment in *Nes.iPKR* amygdala lysates. (n = 3-4 biological replicate lysates, 3-4 mice per group; Two-way ANOVA followed by post-hoc Bonferroni’s test). e) Time course for ciPSI carried out by collecting amygdala lysate at different time points following ASV infusion (0, 0.5h, 1h, 3h and 6h) shows peak expression of 36 kDa iPKR at 0.5h, which was undetectable at 6h (****p<0.0001). Endogenous PKR (68 kDa) remained unchanged after ASV treatment. Peak expression of p-eIF2α, normalized for t- eIF2α, was achieved at 0.5h after ASV infusion followed by a steady decline from 3h onwards (*p<0.05). ATF4 and GADD34 proteins whose transcripts harbor a uORF, were increased by ASV by 1h and and decline to baseline by 6h (***p<0.001 and **p<0.01). cFOS levels significantly declined from baseline at 3h and 6h post infusion (**p<0.01 and ***p<0.001). (n = 3/ group; Repeated measures One-Way ANOVA with post-hoc Bonferroni’s test). f) Schematic for Late-phase long- term potentiation (L-LTP) recording in the lateral amygdala (top) and representative field potentials before (1) and 90 min after tetanus (2) for different groups of slices are shown at the bottom of this panel. L-LTP evoked by 3 trains of high frequency stimulation (HFS) was impaired in Nes.iPKR amygdala slices treated with ASV (5 nM) applied before the tetanus and perfused for 90 min following tetanus (****p<0.0001, n = 12-15 slices/ group). One-way ANOVA with Bonferroni’s post-hoc test. g) Mean fEPSPs at baseline (10 min) before ASV application, at 30 min (i.e. 10 min after ASV application), at 60 min (30 min after tetanus) and at 120 min (90 min after tetanus). ASV significantly reduced fEPSP slope in Nes.iPKR amygdala at 120 min compared to vehicle treatment (***p<0.01) and ASV treated wild-type amygdala (****p<0.001), but had no effect on baseline. Repeated measures Two-way ANOVA with Bonferroni’s post-hoc test. Genotype/Drug X Time interaction: F(6,144) = 4.688, ***p=0.0002; Genotype/Drug: F(2,144) = 171.4, ****p<0.0001; Time: F(3,144) = 5.684, **p=0.0010. Data are presented as mean +/− SEM.

Past studies have shown that the enduring late phase of LTP (L-LTP) in amygdala requires protein synthesis^29, 30^. To determine whether chemogenetic inhibition of pan-neuronal protein synthesis influences synaptic plasticity, we examined long-term potentiation (LTP) of lateral amygdala. We delivered three trains of high-frequency stimulation (HFS) to thalamo-amygdalar inputs and recorded the field excitatory postsynaptic potential (fEPSP) in the dorsal subdivision of lateral amygdala while perfusing into the amygdala slices 5 nM ASV 10 min before and 90 min after tetanus. We found that L-LTP was significantly inhibited by ASV-induced release of iPKR (Fig. 2f). The drug had no effect on baseline fEPSP slope in both wild-type and Nes.iPKR slices (Fig. 2g). The inhibition had a rapid onset after tetanus and became more robust during L-LTP maintenance (Fig. 2g). Together, these results indicate that chemogenetic inhibition of protein synthesis impairs the expression of L-LTP.

### Temporally structured protein synthesis is required for LTM consolidation

Given our novel approach that bypasses the limitations and side effects of methods used to study protein synthesis *in vivo*, we could now study the role of *de novo* protein synthesis in memory consolidation. We trained Nes.iPKR and control mice in simple auditory threat conditioning and centrally infused ASV immediately after training (Fig. 3a). Nes.iPKR and wild-type mice were comparable in learning the association of tone with foot shock, as well as in short-term memory (STM) that is protein synthesis-independent (Fig. 3b and c). However, when the animals were tested for LTM 24h later, Nes.iPKR mice infused with ASV exhibited markedly reduced defensive behavior in response to the paired tone presentations, even though outside of tone presentations the mice had similar motion indices (Fig. 3d, e, and f). Our findings are consistent with previous studies that showed genetic deletion of eIF2α kinases, GCN2 and PKR, lead to an enhancement of long-term spatial and threat memories and decreases the threshold for L-LTP induction^31, 32^. To ensure that ciPSI did not permanently damage the memory system, we retrained Nes.iPKR mice previously infused with ASV in the auditory threat conditioning paradigm. Nes.iPKR mice fully recovered LTM and exhibited a species-appropriate defensive response to the conditioned tone (Fig. 3e and f). Because Nes.iPKR mice express viral NS3/4 protease in addition to the actuator iPKR, we tested the molecular specificity of ciPSI with the PKR inhibitor C16, which binds to PKR at the ATP-binding site thereby blocking its kinase activity^33^. Pre-training administration of C16 inhibited the activation of iPKR and prevented the LTM deficit in Nes.iPKR mice (Fig. 3e and f). Depending on the training paradigm, there are multiple critical periods during memory consolidation that are sensitive to protein synthesis inhibition^9, 34^. Therefore, we assessed whether consolidation of associative threat memory in our paradigm requires one or more waves of *de novo* translation by inducing ciPSI at 0, 1h, and 4h following training (Fig. 3g). Only Nes.iPKR mice receiving ASV infusion immediately after training displayed LTM deficit measured 24h after training (Fig. 3g), suggesting that there is only one wave of protein synthesis for up to 4h following training in our paradigm.

**Figure 3.**
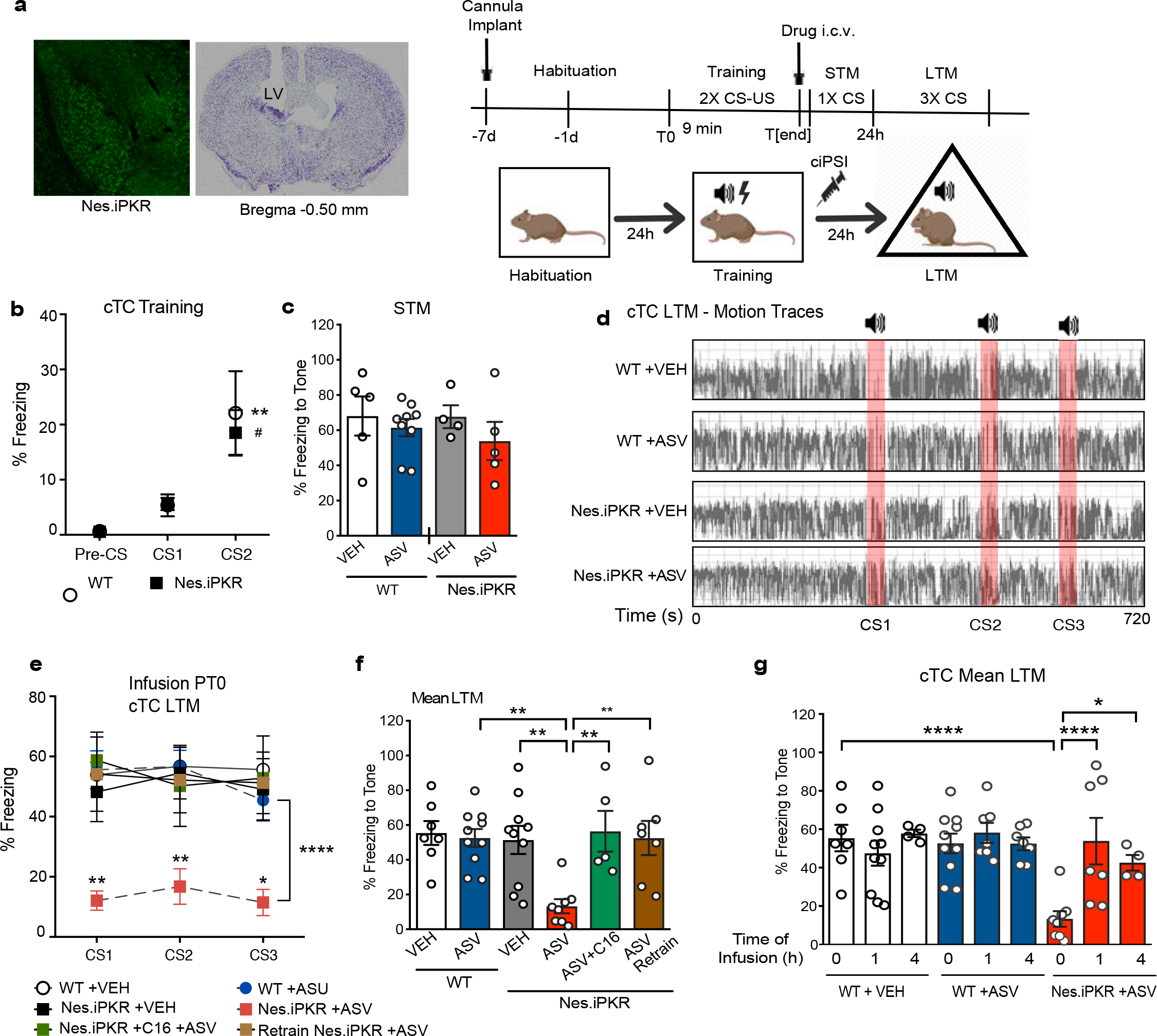
Temporally structured protein synthesis is required for LTM consolidation. a) Schematic of the experimental paradigm for simple cued threat conditioning (cTC) in Nes.iPKR mice pan-neuronally expressing the ciPSI system. b) Nes.iPKR animals learn the association between CS and US (CS2 vs CS1: #p<0.05) similar to the WT mice (**p<0.01) n[WT]=13, n[Nes.iPKR]=16. Repeated Measures Two-way ANOVA with Bonferroni’s post-hoc test. CS: F(2,54) = 19.38, ****p<0.0001. c) Nes.iPKR mice infused with ASV were comparable in STM performance as vehicle-infused mice and WT controls. n=5-9 per group, Two-way ANOVA. d) Representative motion traces for WT mice infused with VEH or ASV, and Nes.iPKR mice infused with VEH or ASV. e) Long-term memory (LTM) tested 24h after training was significantly impaired for Nes.iPKR mice infused with ASV compared to controls for all three CS presentations. Pre-training administration of the PKR inhibitor C16 rescued the memory deficit in Nes.iPKR mice infused with ASV for all three CS presentations. Re-training the Nes.iPKR mice that previously underwent ciPSI-mediated amnesia fully recovers LTM for all three CS presentations p<0.05. n =7-9 per group; Repeated measures Two-way ANOVA with Bonferroni’s post-hoc test. Genotype: F(5,40)=4.570, **p=0.0022. f) cTC Mean LTM was significantly impaired for Nes.iPKR animals treated with ASV compared with VEH (**p=0.021) and WT mice infused with ASV (**p=0.0015). cTC mean LTM in Nes.iPKR mice was rescued with C16 compared with Nes.iPKR mice infused with ASV alone (**p=0.0041). cTC mean LTM2 in retrained Nes.iPKR mice was comparable to WT mice. n=7-10 per group. One-way ANOVA with Bonferroni’s post-hoc test. F(5,41) = 4.890, **p=0.0013. g) Schematic of the experimental paradigm for testing the temporal structure of the requirement for protein synthesis in cTC LTM. h) The LTM deficit was only present for Nes.iPKR mice infused with ASV immediately after training (***p=0.0002) but not 1h or 4h after training. One way ANOVA with Bonferroni’s post-hoc test. n=4-10 per group. Data are presented as mean +/− SEM.

### Cell type-specific *de novo* translation in LA CamK2α-positive neurons is necessary for LTM consolidation

Lesioning and functional inactivation studies have shown that the lateral amygdala (LA) is an integral brain structure for the formation and storage of conditioned aversive memories^4, 5, 35,36^. Learned threat elicits persistent cortical and thalamic input-specific synaptic potentiation in principal excitatory neurons within LA^37, 38^. Thus, we asked whether *de novo* protein synthesis in CamK2α-positive principal neurons in LA is necessary for aversive memory consolidation. We virally delivered CamK2α.Cre.EGFP into the bilateral LA of iPKR mice and control mice (Fig. 4a). Phosphorylation of eIF2α was increased specifically in CamK2α.iPKR neurons compared to wild-type mice and non-EGFP positive neurons in the same animals (Fig. 4b) with a corresponding decrease in *de novo* translation (Fig. 4c) upon ASV administration. CamK2α.iPKR mice were threat conditioned as previously and centrally infused with ASV immediately after training (Fig. 4d). CamK2α.iPKR mice were comparable to wild-type controls in memory acquisition (Fig. 4e), but LTM tested 24h later was significantly impaired (Fig. 4f-h). This LTM consolidation deficit with ciPSI was prevented by pre-training administration of ISRIB, a drug that rescues the p-eIF2α mediated constraint on general translation by enhancing the activity of eIF2B (Fig. 4g-h)^39^. We also tested ASV-treated CamK2α.iPKR mice in open field test and elevated plus maze to gauze anxiety related behaviors and found that although spontaneous locomotion was normal in these animals, there was a significant increase in the percentage of time spent in the open arm, indicating reduced anxiety related behavior (Supplemental Fig. 4a-e).

**Figure 4.**
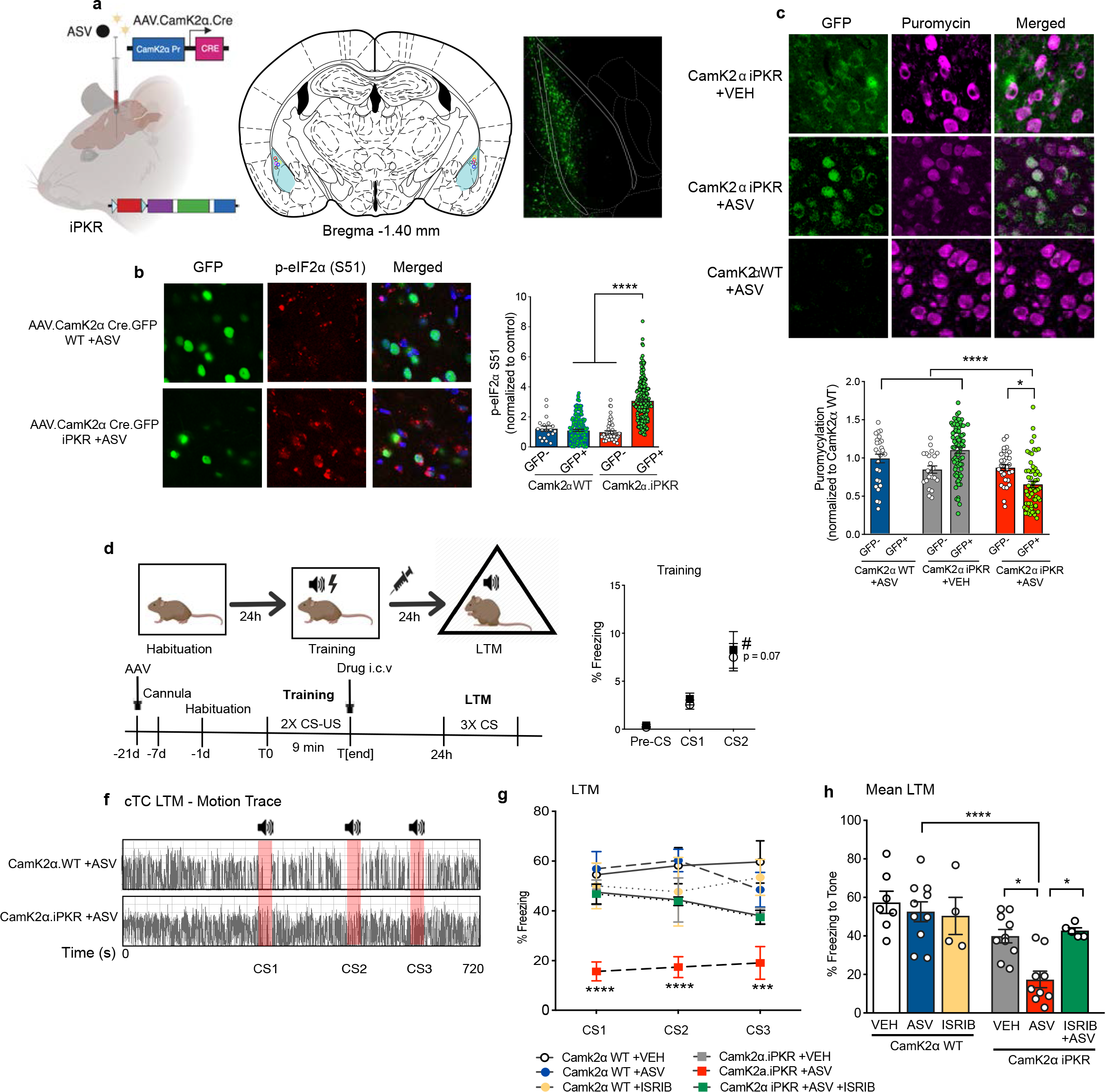
Cell type-specific protein synthesis inhibition in principal neurons in lateral amygdala. a) CamK2α.iPKR mice were generated by bilaterally injecting AAV1.CamK2α.Cre (or AAV9.CamK2α.Cre.EGFP) into the LA of iPKR mice. b) ASV infusion caused a significant increase in p-eIF2α levels in GFP+ neurons in CamK2α.iPKR mice compared to GFP-neighboring neurons in LA (****p<0.0001) as well as GFP+ neurons in CamK2α WT mice (****p<0.0001). Both wild-type and iPKR mice were injected with AAV.CamK2α.Cre.GFP in LA. n CamK2α WT +ASV = 19 (GFP−), 170 (GFP+), n CamK2α.iPKR +ASV = 49 (GFP−), 193 (GFP+) neurons from 3 mice/group. Two-way ANOVA with Bonferroni’s post-hoc test. Genotype X GFP interaction: F(1,427) = 45.11, ****p<0.0001; Genotype: F(1,427) = 28.53, ****p<0.0001; GFP: F(1,427) = 36.75, ****p<0.00001. c) *In* vivo SUnSET assay: ASV treated CamK2α.iPKR mice have significantly reduced translation in LA principal neurons compared to VEH treatment (****p<0.0001) and ASV-treated WT mice (****p<0.0001). One-way ANOVA followed by Bonferroni’s post-hoc test. n CamK2α WT +ASV = 36 (GFP−), n CamK2α.iPKR +VEH = 22 (GFP−), 78 (GFP+) and n CamK2α.iPKR +ASV = 31 (GFP−), 58 (GFP+) neurons from 3 mice/group. d) Schematic of experimental paradigm for cued threat conditioning (cTC) in CamK2α.iPKR mice. e) CamK2α.iPKR mice were comparable with WT in learning the CS-US association. RM Two-way ANOVA with Bonferroni’s post-hoc test. CS: F(2,24) = 24.93, ****p<0.0001 f) Representative cTC LTM motion traces for CamK2α WT and CamK2α.iPKR. g) CamK2α.iPKR mice receiving ASV infusion immediately after training displayed a severe LTM deficit across all three CS presentations (****p<0.0001) that was rescued with pre-training infusion of ISRIB. CamK2α WT +VEH n=7, CamK2α WT +ASV n=9, CamK2α.iPKR +ASV n=9, and CamK2α.iPKR +ASV+ISRIB n=5; RM Two-way ANOVA with Bonferroni’s post-hoc test. Genotype: F(5,35) = 9.767, ****p<0.0001. h) cTC mean LTM was impaired in CamK2α.iPKR mice infused with ASV (****p<0.0001) but was rescued by ISRIB administration (*p<0.05). One-way ANOVA with Bonferroni’s post-hoc test. Data are presented as mean +/− SEM

It is evident that the translation program regulated by eIF2α is necessary for LTM consolidation; however, the accumulation of ATF4 raises the question of whether it is either decreased protein synthesis or increased ATF4 levels causing the memory deficit. Indeed, a previous study has reported that even in the absence of translation inhibition, the phosphorylated eIF2α-mediated increase in ATF4 levels can cause memory deficit by transcriptional repression of CREB regulated genes^15^. Dissociating the effects of general translation inhibition from that of ATF4 expression on memory processes is not trivial because a pharmacological inhibitor of ATF4 does not exist and moreover, long-term knockdown of ATF4 causes deficits in synaptic plasticity and long-term memory^40^. We therefore devised an alternate chemogenetic strategy by blocking translation initiation in LA CamK2α*+* neurons with a knock-in mouse line for expressing tet-dependent synthetic micro-RNA specific for eukaryotic initiation factor 4E (eIF4E) and injected a cocktail of AAV.CamK2α.Cre and AAV.DIO.tTA into the bilateral LA^41^ (Supplemental Fig. 5a) to achieve eIF4E knockdown. eIF4E is a component of the eIF4F complex that binds the 5’ m^7^GTP cap found on majority of cellular mRNAs, and the availability of free eIF4E is tightly regulated in mammalian cells by eIF4E-binding proteins such as 4E-BP and CYFIP1^17^. Knocking down eIF4E in LA CamK2α*+* neurons did not impair learning in the auditory threat conditioning paradigm (Supplemental Fig. 5b), but strongly impaired LTM (Supplemental Fig. 5c and d). Together, the LTM deficit observed using two independent approaches for blocking translation initiation in CamK2α.iPKR and CamK2α.4Ekd respectively strongly support our hypothesis that *de novo* translation in LA Camk2α+ neurons is necessary for consolidation of long-term threat memories.

### Bidirectional control of memory strength by phosphorylation of eIF2α in LA principal neurons

Constitutive heterozygous phospho-mutant eIF2α S51A (A/+) mice have enhanced LTM in several memory paradigms including contextual and auditory threat conditioning and conditioned taste aversion^14^, indicating that phosphorylation of eIF2α is a malleable constraint on protein synthesis during memory consolidation. Therefore, we virally delivered Camk2α.Cre.EGFP bilaterally to the LA of floxed phosphomutant eIF2α (S51A) mice^42^ (Fig. 5a). Targeting phosphomutant eIF2α S51A expression in LA CamK2α+ neurons led to a significant increase in *de novo* translation in homozygous CamK2α.eIF2α (A/A) but not in heterozygous CamK2α.eIF2α (A/+) neurons, as assessed with *in vivo* SUnSET (Fig. 5b), and also led to a decrease in phosphorylated eIF2α (Supplemental Fig. 6a). Congruently, although all of the mice learned auditory threat conditioning (Fig. 5c), homozygous CamK2α.eIF2α (A/A) mice exhibited significantly higher freezing responses to the conditioned tone in the LTM test, indicating more robust memory compared to wild-type controls (Fig. 5e-f). We next sought to correct the memory deficit in the CamK2α◻iPKR mice by introducing phosphomutant eIF2α, and thus generated CamK2α.iPKR eIF2α (A/+) and CamK2α.iPKR eIF2α (A/A) animals by injecting AAV.CamK2α.Cre in the LA of mice double transgenic for iPKR and eIF2α S51A. Memory deficits in the CamK2a.iPKR mice were completely rescued by dephosphorylating both alleles of eIF2α, but was unaltered by dephosphorylating one allele of eIF2α (Fig. 5e-f). This indicated that the translation tone and LTM consolidation process is set by the phosphorylation of eIF2α, even when the abundance of phosphorylatable eIF2α is reduced by half, and confirmed that the memory deficit caused by iPKR is through the phosphorylation of eIF2α and not due to non-specific cellular toxicity.

**Figure 5.**
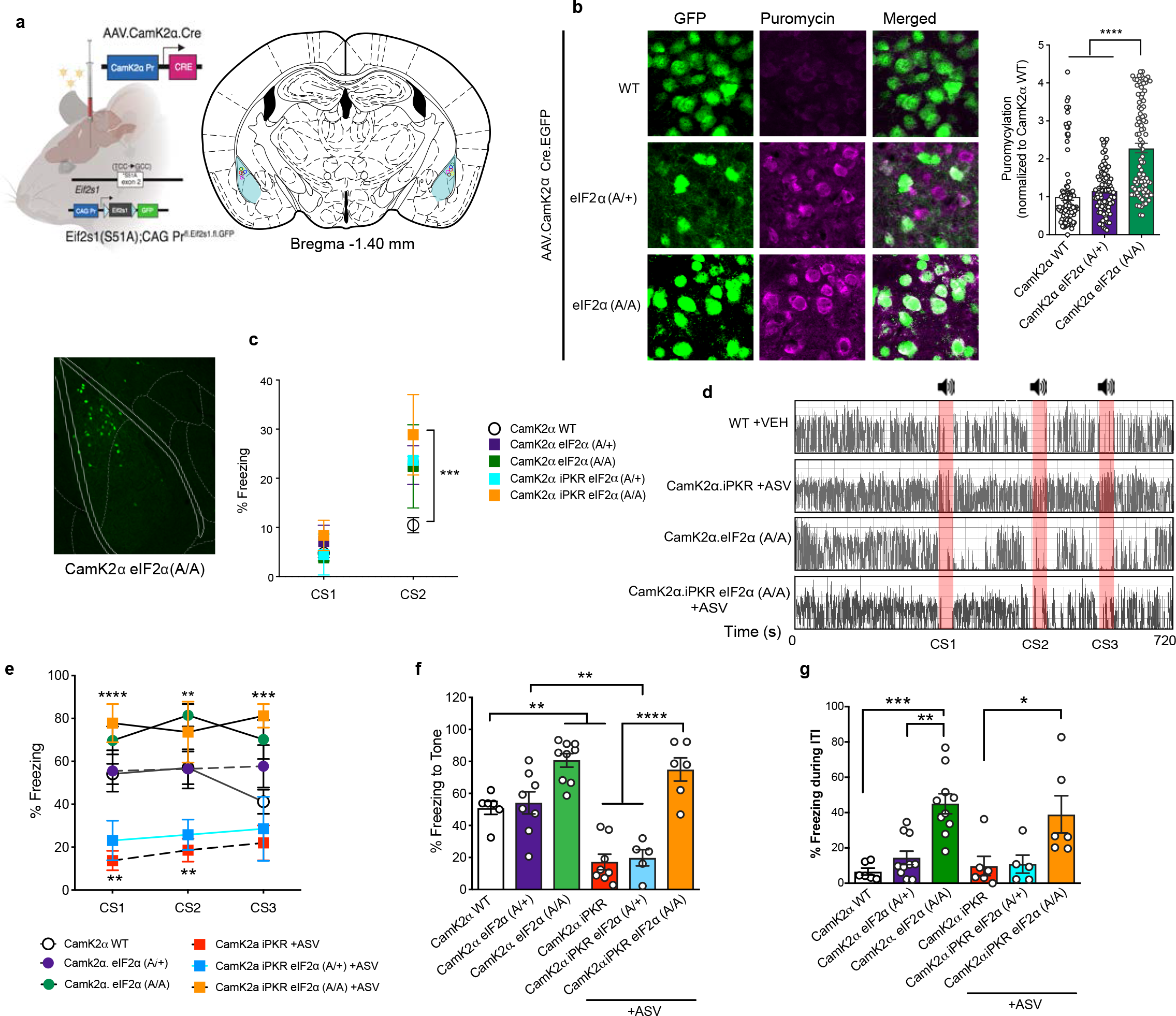
Bidirectional control of memory strength by phosphorylation of eIF2α in LA principal neurons. a) Heterozygous CamK2α.eIF2α (A/+) and homozygous CamK2α.eIF2α (A/A) were generated by bilaterally injecting AAV1.CamK2α.Cre into the lateral amygdala of eIF2α (S51A) CAG Pr^.fl.eIF2α.fl.GFP^ mice. b) *In vivo* SUnSET assay: Homozygous CamK2α.eIF2α (A/A) mice displayed a significant increase in *de novo* translation compared to the CamK2α wild-type mice and heterozygous CamK2α.eIF2α (A/+) mice (***p<0.001). n = 93-101 neurons (from 3 mice) per group; One-way ANOVA followed by Bonferroni’s post-hoc test. F(2,284) = 52.01, ****p<0.0001. c) Double transgenic iPKR eIF2α (A/+) and iPKR eIF2α (A/A) mice were bilaterally injected with AAV1.CamK2α.Cre in the lateral amygdala to generate CamK2α.iPKR eIF2α (A/+) and CamK2α.iPKR eIF2α (A/A) mice, respectively. All groups of mice (CamK2α WT, CamK2α eIF2α (A/+), CamK2α.eIF2α (A/A), CamK2α.iPKR eIF2α (A/+) and CamK2α. iPKR eIF2α (A/A)) learned the CS-US association, however CamK2α.eIF2α (A/A) and CamK2α. iPKR eIF2α (A/A) mice learned better than CamK2α WT mice (CS2: CamK2α WT vs CamK2α. iPKR eIF2α (A/A), ***p=0.0009; CamK2α WT vs CamK2α.eIF2α (A/A), p = 0.0740). RM Two-way ANOVA with Bonferrroni’s post-hoc test. CS: F(1,31) = 44.25, ****p<0.0001. d) Freeze frame motion traces for CamK2α WT mice treated with VEH, CamK2α iPKR mice treated with ASV, and CamK2α eIF2α (A/A) mice. e-f) In the cTC LTM test, homozgygous CamK2α.eIF2α (A/A) mice displayed enhanced memory compared to CamK2α WT mice (**p<0.01), whereas as shown previously, CamK2α.iPKR mice treated with ASV have impaired memory (**p<0.01). Heterozygous CamK2α eIF2α (A/+) mice displayed comparable LTM as WT mice. Heterozygous CamK2α.iPKR eIF2α (A/+) mice treated with ASV exhibited comparable memory deficit as CamK2α iPKR mice treated with ASV. However, the memory deficit in CamK2α.iPKR mice was fully rescued by introducing non-phosphorylatable eIF2α in homozygous CamK2α.iPKR eIF2α (A/A) mice (****, p<0.0001). n = 6-9 per group; RM Two-way ANOVA with Bonferroni’s post-hoc test. Genotype: F(5,35) = 12.79, ****p<0.0001. f) Bar graphs for mean cTC LTM for all groups. One-way ANOVA with Bonferroni’s post-hoc test. F(5,36) = 23.30, ****p<0.0001. g) Memory enhancement in CamK2α eIF2α (A/A) mice was accompanied by cognitive inflexibility at the offset of CS. During intertrial intervals (ITI), CamK2α eIF2α (A/A) freeze at a much higher rate compared to the WT (***p<0.001) and CamK2α eIF2α (A/+) mice (**p<0.01). Similarly, CamK2α.iPKR eIF2α (A/A) +ASV mice displayed a higher freezing rate during the ITIs compared to CamK2α iPKR treated with ASV (*p<0.05). n=6-9 per group; One-way ANOVA with Bonferroni’s post-hoc test. F(5,36) = 8.569, ****p<0.0001. Data are presented as mean +/− SEM

Notably, the memory enhancement in the CamK2α.eIF2α (A/A) mice came with a cost. We found that the dysregulated increase in translation tone in CamK2α.eIF2α (A/A) mice resulted in impaired behavioral flexibility during the tone offset in LTM test. CamK2α.eIF2α (A/A) mice freeze starting at tone onset, but continue to freeze after the tone offset during inter-trial interval epochs (Supplemental Fig. 6b and Fig. 5g). This failure to switch off defensive state is triggered only after the onset of first tone while at pre-conditioned stimulus (CS) the animals have normal motor behavior, indicating absence of generalized anxiety or contextual threat response. We next tested CamK2α.eIF2α (A/A) animals in open field arena and elevated plus maze, and found normal locomotion in open field (Supplemental Fig. 6c-e), but a decrease in the percentage of time spent in the open arm in the elevated plus maze, indicative of increased anxiety-like behavior (Supplemental Fig. 6f-h). These data suggest that a dysregulated increase in the translation program regulated by eIF2α leads to a failure in risk assessment in non-threatful environments and epochs.

### Dysregulated protein synthesis in LA CamK2α+ neurons compromises the precision of a complex memory

Finally, we addressed whether lost memories can be rescued with artificial reactivation^43^ of LA CamK2α+ principal neurons. Designer drug activated by designer drug (DREADD) hM3Dq mediated neuronal activation engages the mitogen activated protein kinases, ERK1/2, and mTORC1 pathways that are positively associated with protein synthesis^44, 45^. We injected a cocktail of AAV.CamK2α.Cre together with AAV.hSyn.DIO.hM3Dq.mCherry into the LA of iPKR knock-in mice (Fig. 6a). DREADD agonist C21^46^ significantly increased *de novo* translation in CamK2α+ neurons of CamK2α.iPKR hM3Dq mice compared to vehicle treated group (Fig. 6b). We then trained CamK2α.iPKR hM3Dq, CamK2α.hM3Dq, and wild-type control mice in a differential threat conditioning paradigm that involves three interleaved presentations of a paired tone (CS+) and an unpaired tone (CS−) in a single session (Fig. 6c-d). Consistent with earlier data, post-training ASV infusion impaired LTM and led to a significant decline in freezing response to paired tone (Fig. 6f-g). 48h after LTM1 test, we re-tested the animals for LTM2 in a different context following chemogenetic activation of hM3Dq receptors in the CamK2α*+* neurons using agonist C21. We found that although artificial reactivation of CamK2α+ neurons recovered CS+ LTM, it led to stimulus generalization and resulted in generalized defensive freezing response to CS− (Fig. 6h) that reflected in a significant decline in threat discrimination index (Fig. 6i). CamK2α.wild-type and CamK2α.hM3Dq mice exhibited no change in discrimination index between the two LTM tests. Besides stimulus generalization, freezing during ITI was also increased by artificially boosting protein synthesis in translation-inhibited CamK2α+ neurons (Supplemental Fig. 7a-b).

**Figure 6.**
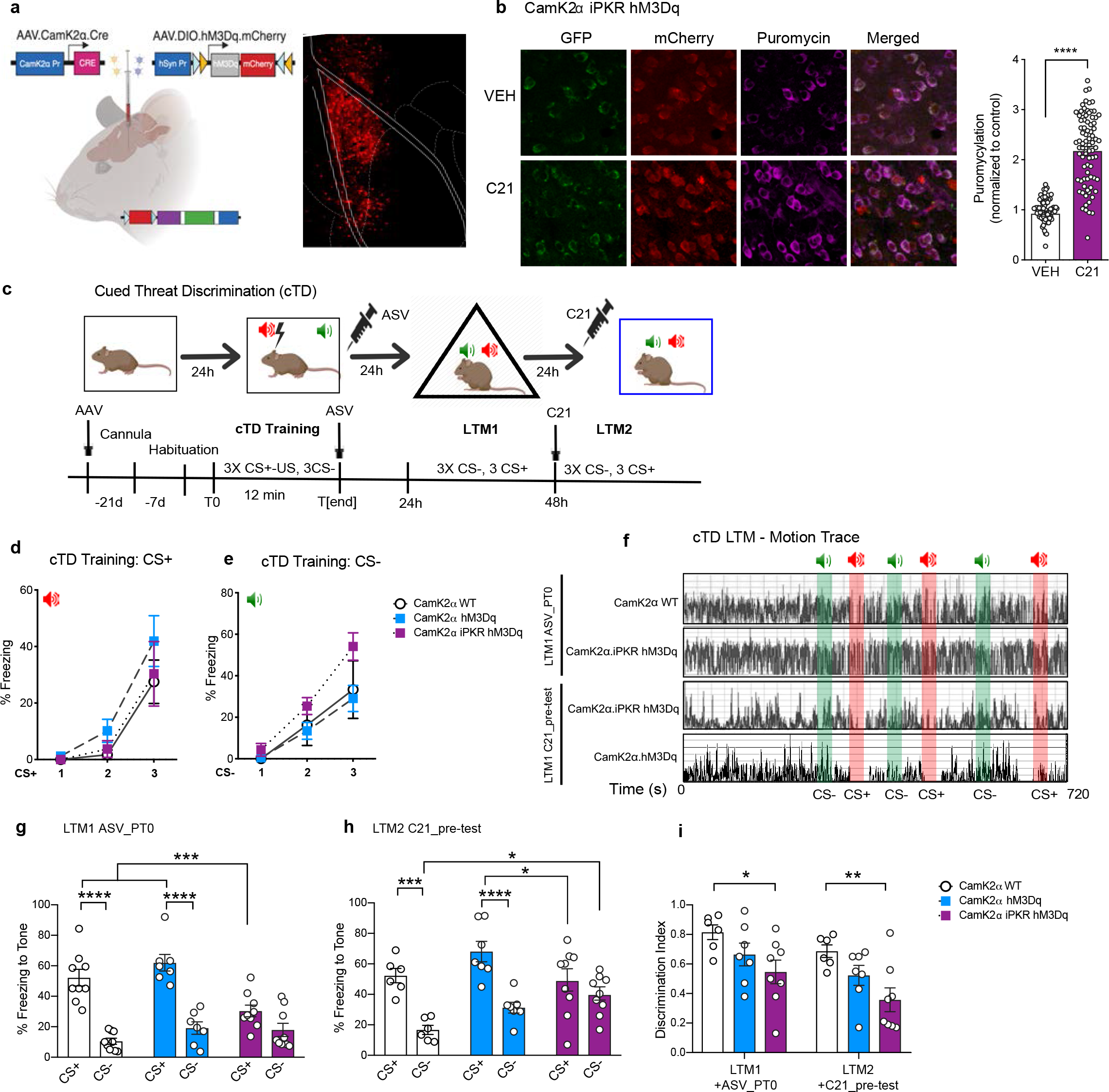
Artificial reactivation of LA principal neurons compromises the precision of a complex memory. a) CamK2α.iPKR.hM3Dq and CamK2α.hM3Dq mice were generated by bilaterally injecting AAV1.CamK2a.Cre and AAV9.hSyn.DIO.hM3Dq.mCherry into the LA of either iPKR or wild-type mice. b) DREADD receptor agonist C21 significantly enhanced *de novo* translation, as measured with *in vivo* SUnSET, in CamK2α.iPKR.hM3Dq neurons compared to vehicle treatment (****p<0.0001). Unpaired t-test. n = 74−82 neurons from 3 mice per group. c) Schematic of experimental paradigm for cued threat discrimination conditioning (cTD) in CamK2α.iPKR.hM3Dq mice. d) CamK2α WT, CamK2α.hM3Dq, and CamK2α.iPKR hM3Dq mice learned equivalently to associate CS+ with US during training, although they did not yet develop discrimination of CS− from CS+ during training (e). f) Representative cTD LTM motion traces for WT mice, CamK2α.iPKR.hM3Dq mice that were infused with ASV immediately after training, CamK2α.iPKR.hM3Dq mice that underwent reactivation of LA principal neurons with C21 and CamK2α. eIF2α (A/A) mice. g) CamK2α.iPKR.hM3Dq mice infused with ASV immediately after training exhibited a significant CS+ memory deficit compared to ASV treated WT CamK2α.hM3Dq mice (**p<0.01). One-way ANOVA with Bonferroni’s post-hoc test. Genotype/Drug X CS interaction: F(2,44) = 8.30, ***p=0.0009; Genotype/Drug: F(1,44) = 83.69, **p=0.0025; CS: F(2,44) = 6.904, ****p<0.0001. h) Artificial reactivation of LA principal neurons in CamK2α.iPKR.hM3Dq neurons using C21 recovered CS+ LTM (**p<0.01), but caused stimulus generalization to CS− (*p<0.05). One-way ANOVA with Bonferroni’s post-hoc test. Genotype/drug X CS interaction: F(2,19) = 6.888, **p=0.0056; CS: F(1,19) = 55.84, ****p<0.0001. f) cTD discrimination index was significantly decreased in CamK2α.iPKR.hM3Dq mice that under artificial reactivation following ciPSI compared to CamK2α.hM3Dq WT mice (*p<0.05), as well as CamK2α.iPKR.hM3Dq mice following ciPSI (*p<0.05). One-way ANOVA with Bonferroni’s post-hoc test. Genotype: F(2,36) = 8.604, ***p=0.0009, LTM test: F(1,36) = 6.773, *p=0.0134. n=6-10 mice per group. Data are presented as mean +/− SEM

To further understand what happens when eIF2α-controlled general translation in LA principal neurons is boosted during consolidation of a complex memory, we explored differential threat conditioning in CamK2α eIF2α S51A mice (Supplemental Fig. 7c). We found that whereas homozygous CamK2α.eIF2α (A/A) mice can discriminate between CS+ and CS− during LTM, they have a significantly increased freezing response to CS−. On the other hand, CamK2α wild-type mice robustly discriminated CS+ from CS- and displayed negligible freezing response to CS− (Supplemental Fig. 7d). The enhanced CS− response in theCamK2α.eIF2α (A/A) mice resulted in a poor discrimination index (Supplemental Fig. 7e). We next examined the freezing during ITI, and found that similar to earlier results with simple threat conditioning, CamK2α.eIF2α (A/A) exhibited a significantly increased freezing response following tone offset, indicating behavioral inflexibility (Supplemental Fig. 7f-g). These findings indicate that the precision of a memory trace is contributed by the finely regulated translation program in LA CamK2α+ neurons during memory consolidation.

## Discussion

Protein synthesis is metabolically expensive and thus is tightly regulated at the level of initiation but until now, an effective chemogenetic tool to block protein synthesis has been lacking, which has limited investigation of cell autonomous protein synthesis in physiological processes. To address this issue, we have bioengineered a spatiotemporally precise chemogenetic resource for rapidly and reversibly blocking cell autonomous protein synthesis via phosphorylation of eIF2α. Phosphorylation of eIF2α is a tightly regulated molecular event that acts as a master effector of the integrated stress response. Using our chemogenetic iPKR mouse resource, we made several notable discoveries: 1) Temporally structured pan-neuronal protein synthesis is required for long-term memory consolidation. Recent long-term memory examined 24 hours after training is most sensitive to protein synthesis disruption in the first hour after training; 2) Blocking *de novo* translation in CamK2α+ principal neurons within lateral amygdala disrupts long-term memory consolidation. This block of memory consolidation can be rescued by either using the eIF2B activator ISRIB or by dephosphorylating both alleles of eIF2α; 3) Expressing a biallelic phosphomutant eIF2α in CamK2α+ principal neurons results in enhanced strength of the memory, but introduces behavioral inflexibility and a generalized defensive response for an unpaired or safe tone; 4) Artificial reactivation of LA CamK2α+ neurons 24h after protein synthesis inhibition recovers lost memory for the paired tone but causes stimulus generalization.

### Protein synthesis dependence during memory consolidation

We found that too little or too much of phosphorylation of eIF2α causes aberrant memory storage and expression. A parsimonious interpretation of our data is that state-dependent modulation of eIF2α phosphorylation is critical for a finely tuned translation program supporting the associative memory. eIF2α phosphorylation itself is a homeostatic process wherein the phosphorylation event triggers the synthesis of the eIF2α dephosphorylating enzyme GADD34, bringing the system back to a lower state of eIF2α phosphorylation. Our data is consistent with Batista *et al* (2016) showing that phosphorylation of eIF2α causes a block of general translation^47^. In contrast, Jiang et al (2010)^15^ reported that chemogenetic dimerization of FKBP-PKR (fPKR) induced phosphorylation eIF2α by 1.5 fold without blocking general translation and thus, they attributed the memory deficit in fPKR expressing animals to ATF4 expression. It is possible that because these investigators injected the drug inducer of fPKR (AP20187) 2h before radioisotope S^35^ methionine injection i.p., the sensitive time window for protein synthesis disruption was missed during labeling. Compared to fPKR, PKR kinase domain (PKRk) is more effective in suppressing translation because it not only directly phosphorylates eIF2α, but it also interacts with and phosphorylates endogenous full-length PKR, further boosting eIF2α phosphorylation^48^. In our system, central infusion of ASV resulted in 2-fold increase in phosphorylated eIF2α, which resulted in a robust 50% decrease of general translation as assessed with *in vivo* SUnSET that involved local infusion of puromycin directly into lateral amygdala. Our convergent data from a parallel strategy of cell autonomous protein synthesis inhibition in LA CamK2α+ neurons using a synthetic micro-RNA against eIF4E provides further support for the requirement of *de novo* protein synthesis in long-term memory consolidation. Nonetheless, we also observed an increase in ATF4 levels following phosphorylation of eIF2α and cannot exclude the possibility that the memory deficit in the iPKR transgenic mice is in part due to upregulated translation of transcripts containing uORFs, such as ATF4 and the ensuing repression of CREB-regulated genes.

### Memory generalization and behavioral inflexibility as a result of dysregulated translation

Several studies have used IEG-based engram cell targeting approaches to interrogate the nature of memory formation and recall. Artificial reactivation of conditioned threat engram cells in LA using optogenetics has been shown to recover memories previously lost by protein synthesis inhibition using anisomycin^49, 50^. Our chemogenetic iPKR mouse resource is not amenable for the engram cell targeting approach because the engram cells by definition need to be tagged during training event for iPKR expression, a process dependent on *de novo* transcription and translation. Nonetheless, our results show that artificial reactivation of previously translation-inhibited CamK2α+ neurons in LA results in memory recovery, but at the cost of stimulus generalization. We propose that learning-induced somatic and synaptic protein synthesis functions to stabilize the connections between the pathway-specific afferents from conditioned tone processing brain regions (auditory cortex and auditory thalamus) to amygdalar engram cells in such a way that the conditioned tone can access and activate the downstream neuronal network in the LA during recall. This natural cue-elicited neuronal reactivation is incompletely recapitulated by artificial reactivation of LA neurons, such that there is imprecise restoration of the memory, perhaps due to a lack of reinstatement of synapse specificity of the original memory trace. Phospho-mutant eIF2α mice, with a rigid increase in translation in LA principal neurons, also exhibit stimulus generalization and behavioral inflexibility at tone offset. Taken together, our data indicate that a finely tuned translation program in the amygdala is required to coordinate the stability and precision of the long-term memory trace.

### Model of protein synthesis regulation during long-term memory consolidation

Based on our data, we propose the following cellular/molecular model of protein synthesis regulation in lateral amygdala during long-term consolidation. Cued threat conditioning leads to a coordinated increase in protein synthesis in CamK2α+ neurons in the soma and in local subcellular compartments, such as dendritic spines and axons. This results in robust memory strength and precision (Fig. 7a). Chemogenetic inhibition of protein synthesis using iPKR blocks general translation in LA CamK2α+ neurons resulting in memory loss (Fig. 7b). Expression of biallelic phosphomutant eIF2α in LA CamK2α+ neurons, although insensitive to iPKR-mediated eIF2α phosphorylation, results in an aberrant increase in basal translation. This leads to robust memory strength, but low memory precision. (Fig. 7c). Artificial chemogenetic reactivation of CamK2α+ neurons following inhibition of protein synthesis leads to an increase in protein synthesis, but does not recapitulate learning-induced local translation, thus causing memory generalization. This is manifested as normalized memory strength and low memory precision (Fig. 7d).

We have used multi-pronged chemogenetic approaches to investigate the translational control of long-term threat memories. Future studies will be required to elucidate how protein synthesis regulation occurs in genetically defined and functionally coherent cell types within the memory network during long-term memory consolidation.

**Figure 7.**
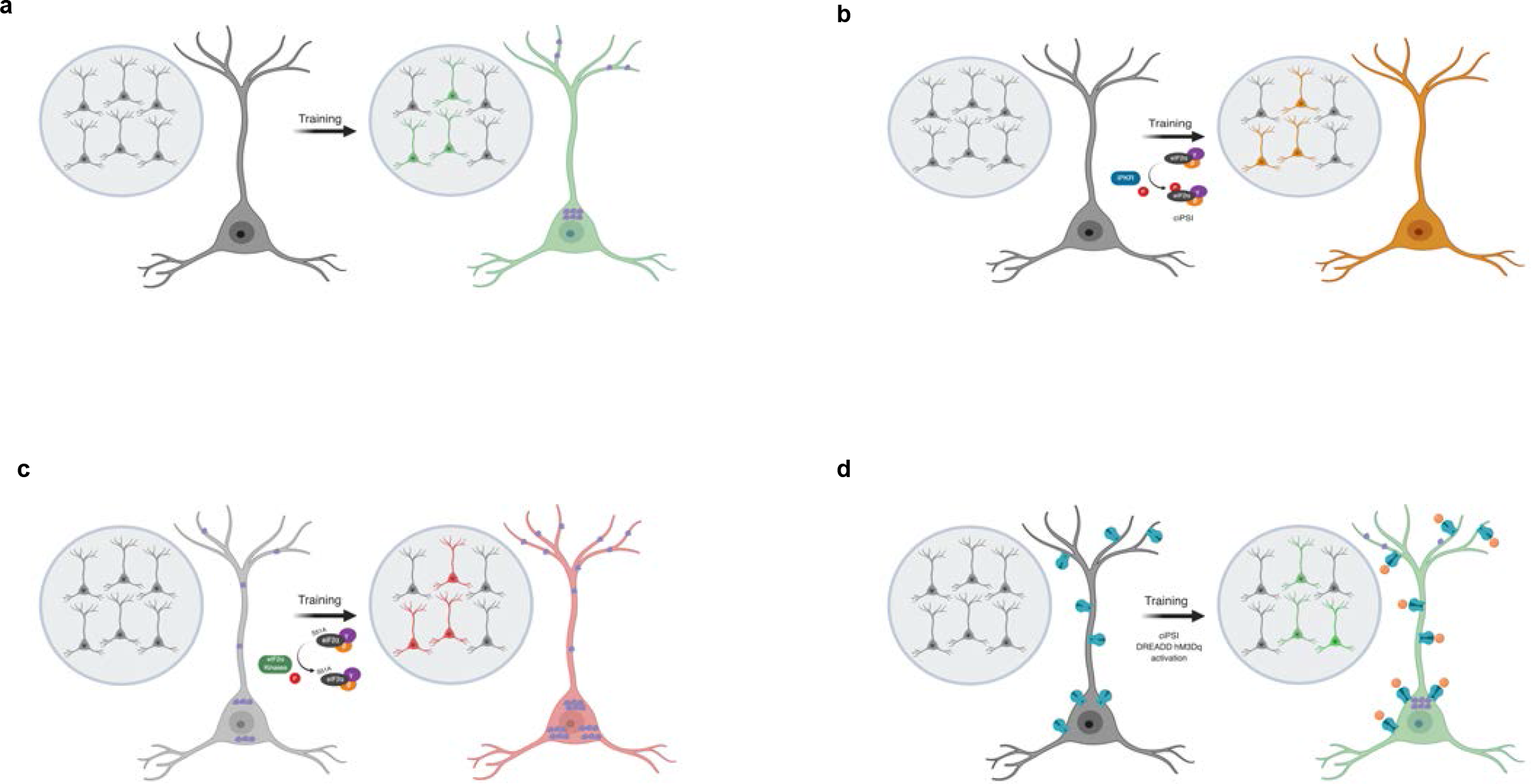
Model for protein synthesis regulation during long-term memory consolidation. a) In wild-type mice, threat conditioning leads to a spatiotemporally regulated increase in somatic and dendritic protein synthesis that stabilizes the memory trace. b) Application of ciPSI system prevents the coordinated increase in cell-wide translation leading to impaired LTM. c) Dephosphorylation of eIF2α enhances general translation, but it is dysregulated and unable to coordinate the cell-wide translation program to store a complex memory trace, resulting in memory generalization. d) Artificial reactivation of the amygdala principal neurons after protein synthesis inhibition-mediated amnesia leads to an increase in translation but does not restore synapse specificity and thus leads to memory generalization.

## ONLINE METHODS

### Cloning of recombinant plasmids

iPKR was produced by adding NS5A/B junction sequence (DEMEECASHLPY) within the PKR kinase domain. NS3/4 protease, iPKR, and GFP were amplified using Phusion polymerase (NEB) and cloned into peGFP-N1 vector (Clontech) using appropriate restriction enzymes (NEB). All constructs were verified by sequencing. Purifications were done using PCR extraction kit (Qiagen). The ligations were performed with Quick Ligase kit (NEB) and the products were transformed into chemically competent TOP10 cells (Invitrogen) that were grown in LB medium containing 35 mg/ml Kanamcyin (Sigma). The mammalian expression constructs were sub-cloned into the gene targeting plasmids as previously reported^31^. The final construct was extracted with phenol: chloroform: isoamyl alcohol (25:24:1), precipitated in 70% ethanol and dissolved in Tris-EDTA buffer.

### Cell culture and transfection

293T cells were grown in a 24-well plate in growth medium (Invitrogen) supplemented with 10% fetal bovine serum (Invitrogen) to have 90% confluency on the day of transfection. Transfection was carried out with 1 μg of a reference plasmid using 2 μL 293Tfectin reagent (Invitrogen). The amounts of the remaining plasmids were adjusted to have the same molarity per well. Cells were lysed on ice in phosphate-buffered saline supplied with 1% Triton X-100 (Sigma) and inhibitors against proteases (Pierce, 78437) phosphatases (Pierce, 78420) and translation machinery (100 μg/ml Cycloheximide, Sigma); and the cytosolic extracts were isolated after precipitating the insoluble fraction. The protein concentrations were measured using BCA protein assay (Pierce).

### Generation of iPKR knock-in animals

129S6/SvEvTac embryonic stem cells of W4 parental cell line were used for homologous recombination at *Eef1a1* genomic locus. Gene targeting, generation of knockin mice and Southern blotting was conducted by the Gene Targeting Facility at The Rockefeller University, New York, NY as previously described^14^. Following primers were used for genotyping - β actin forward: GGC TGT ATT CCC CTC CAT CG, β actin reverse: CCA GTT GGT AAC AAT GCC ATG T, EGFP forward: GCA GAA GAA CGG CAT CAA GGT, EGFP reverse: ACG AAC TCC AGC AGG ACC ATG, iPKR forward: CAC CAT GGG CAA GCC CCA GCG TCT GTA TG and iPKR reverse: TGC CGG TCC AGG TGT AGC TCA TGC TGC AGC.

### Animals

Mice were provided with food and water *ad libitum* and were maintained in a 12h/12h light/dark cycle at New York University at stable temperature (78°F) and humidity (40 to 50%). All mice were backcrossed to C57Bl/6J strain for at least 5 generations. Nestin Cre transgenic mice (stock #003771) were obtained from Jackson laboratory as previously described^34^. Nestin Cre mice were bred to floxed iPKR knockin mouse line to generate transheterozygote Nes.iPKR mice. Transgenic homozygous Eif2s1 (S51A); CAG Pr^fl.Eif2s1.fl.GFP^ mice (i.e. eIF2α (A/A)) were a gift from Dr. Randal Kaufman. Double transgenic floxed iPKR/ eIF2α (A/A) or (A/+) mice were obtained by breeding floxed iPKR mice with eIF2α (A/A) mice. Col1a1^TRE GFP.shmir4E.389^ mice were a gift from Dr. Jerry Pelletier. Both male and female mice were used for all experiments. Mice tested in behavioral assays were 10-15 weeks old. All procedures involving the use of animals were performed in accordance with the guidelines of the National Institutes of Health (NIH) and were approved by the University Animal Welfare Committee of New York University.

### Drugs and chemicals

Asunaprevir (ChemExpress) was dissolved in DMSO for a stock concentration of 10 mM and diluted in sterile saline to 100 nM. 2 μl of this drug was intracranially infused into the left lateral ventricle (−0.50 mm anterioposterior [AP], −1,10 mm mediolateral [ML] and −2.20 mm dorsoventral [DV]) using an injection cannula inserted into the stainless steel guide cannula (Plastics One). Except for the dosage curve experiments, 100 nM ASV was used in all experiments involving protein synthesis inhibition *in vivo*. ASV infusion was carried out at 0.5 μl/min using an injection cannula extending out of PE50 tubing attached to a 5 μl Hamilton syringe (Hamilton) using a PHD 2000 Infusion Pump (Harvard Apparatus). After injection, the injection cannula was kept in place for 1 min before its withdrawal. Puromycin (Sigma, P8833) was dissolved in ddH_2_O at 25 μg/μl, and this stock was freshly diluted in saline for SUnSET assays *in vivo.* For SUnSET immunoblot, 2 μl of 25 μg/μl puromycin was infused into the lateral ventricle. For SUnSET immunohistochemistry, 0.5 μl of 10 μg/μl puromycin was infused into the lateral amygdala (−1.40 mm AP, −3.50 mm ML and −4.60 mm DV) of awake behaving mice using internal cannula with 1 mm projection. For BONCAT *ex vivo* assays, azidohomoalanie (AHA) (Fisher, NC0667352) was dissolved in distilled water at 100 mM. 1 mM AHA was bath applied to amygdala slices in ACSF (125 mM NaCl, 2.5 mM KCl, 1.25 mM NaH_2_PO_4_, 25 mM NaHCO_3_, 25 mM glucose, 1 mM MgCl_2_ and 2 mM CaCl_2_). DREADD actuator, agonist C21 (Tocris), was dissolved in DMSO at 40 mg/ml concentration and freshly diluted in saline and administered to mice at 1 mg/kg i.p. 25 mM PKR inhibitor C16 (Cal-biochem) was dissolved in DMSO and diluted in saline to a final DMSO concentration of 0.5%. PKR inhibitor C16 was administered i.p. at a dosage 5 mg/kg 30 min before training when specified. ISRIB was dissolved in 1:1 PEG400:DMSO and injected in mice at 2.5 mg/kg i.p as previously described^33^. Doxycycline was prepared in rodent chow (Bio-Serv) at 40 mg/kg and this diet was provided to the mice in 4Ekd group for two weeks following surgery.

### Stereotaxic surgeries

Mice were anesthetized with the mixture of ketamine (100 mg/kg) and xylazine (10 mg/kg) in saline (i.p. injection). To generate CamK2α.iPKR mice designated for behavior testing, 100 nl of AAV1.Camk2α.Cre (1.6 × 10^13 GC/ml; UPenn Vector Core) was injected bilaterally into the lateral amygdala (−1.40 mm AP, +/−3.50 mm ML, −4.60 mm DV) of iPKR mice. Two weeks after viral injections, CamK2α.iPKR mice underwent a second surgery for implantation of 23 gauge stainless steel guide cannulae (Plastics One) in the left lateral ventricle (−0.50 mm AP, −1.10 mm ML and −2.20 DV). One skull screw was inserted into the skull, and dental cement was applied to secure the cannula in position (Parkell and Stoelting). For *in vivo* SUnSET immunohistochemistry, CamK2α.iPKR mice were implanted with two guide cannulas, one in LV (−0.50 mm AP, −1.10 mm ML and −2.20 DV) for ASV infusion and a second one in LA (−1.40mm AP, +3.50 mm ML and −3.60 mm DV) for puromycin infusion. For assessing phosphorylation of eIF2α on S51, a separate group of CamK2α GFP.iPKR mice were generated by injecting AAV9. Camk2α.Cre.EGFP and AAV9.DIO.tTA in the lateral amygdala. For eIF4E knockdown experiments, 100 nl each of AAV1. Camk2α.Cre and AAV9.CAG.DIO.tTA (1 × 10^13 GC/ml; Vigene) was injected into the LA of Col1a1^TRE GFP.shmir4E.389^ mice. Control wildtype littermates were injected with 100 nl each of AAV1. Camk2α.Cre and AAV9.CAG.DIO.GFP (3.33 × 10^13 GC/ml; UPenn Vector Core). For artificial reactivation studies involving CamK2α.iPKR hM3Dq mice, a mixture of 100 nl each of AAV1.Camk2α.Cre and AAV.hSyn Pr.DIO.hM3Dq.mCherry (2.31 × 10^12 GC/ml; Addgene) were injected bilaterally into the LA of iPKR mice. To generate CamK2α .eIF2α (A/+) and CamK2α.eIF2α (A/A) mice, 100 nl of AAV1.Camk2α.Cre (UPenn vector core) was injected bilaterally into the lateral amygdala (−1.40 mm AP, +/− 3.50 mm ML, −4.60 mm DV) of eIF2α (A/+) or eIF2α (A/A) mice, respectively. Finally, CamK2α.iPKR eIF2α (A/+) and (A/A) mice were generated by injecting 100 nl of AAV1.Camk2α.CRE into the lateral amygdala (−1.40 mm AP, +/−3.50 mm ML, −4.60 mm DV) of double transgenic floxed iPKR and eIF2α (A/+) or eIF2α (A/A) mice, respectively. Control wild-type littermates were injected with AAV9.Camk2α.Cre.EGFP.

### Electrophysiology

The electrophysiology experiments were performed as previously described^29, 51^ and mice were between 2 and 4 months of age at the time of the experiments. Briefly, transverse slices (300 μm) containing the amygdala were isolated and transferred to recording chambers (preheated to 32°C), where they were superfused with oxygenated aCSF (composition in mM: 125 NaCl, 2.5 KCl, 1.25 NaH_2_PO_4_, 25 NaHCO_3_, 25 D-glucose, 2 CaCl_2_ and 1 MgCl_2_) at least 1 hr before recordings began at a rate of 2 ml/min. In most of the experiments picrotoxin (75 μM) was present in the perfusion solution. For bath application the drugs were made and stored as concentrated stock solutions and diluted 1000-fold when applied to the perfusate. Field excitatory postsynaptic potentials (fEPSPs) from the internal capsule pathway were recorded using microelectrodes filled with artificial cerebrospinal fluid (aCSF; resistance 1-4 MΩ). A bipolar Teflon-coated platinum electrode was placed in the thalamic afferent fiber to the lateral amygdala, which is located in the ventral part of the striatum just above the central nucleus of the amygdala. The test stimuli for basal synaptic response was at 0.05 Hz. In all experiments, basal field excitatory postsynaptic potentials (fEPSPs) were stable for at least 20 min before the start of each experiment. L-LTP was induced with three 1 sec 100-Hz high-frequency stimulation (HFS) trains, with an intertrain interval of 60 sec. After induction of L-LTP, we collected fEPSPs for an additional 120 min. 5 nM ASV or vehicle were bath applied 10 min before L-LTP induction and lasted for all the recording. Slope values were compared from the Nes.iPKR transgenic mice and their control littermates treated with either ASV or vehicle. Synaptic efficacy was monitored at 0.05 Hz and averaged every 2 min. fEPSPs were amplified and digitized using the A-M Systems Model 1800 and Digidata 1440 (Molecular Devices)

### Behavior

#### Open field activity

All behavior sessions were conducted during the light cycle. Mice were placed in the center of an open field (27.31 × 27.31 × 20.32 cm) for 15 min during which a computer-operated optical system (Activity monitor software, Med Associates) monitored the spontaneous movement of the animals as they explored the arena. The parameters tested were: distance traveled and the ratio of center to total distance at three epochs.

#### Elevated plus maze

The plus maze consisted of two open arms (30 cm × 5 cm) and two enclosed arms of the same size with 14-cm high sidewalls and an endwall. The arms extended from a common central square (5 cm^2^ × 5 cm^2^) perpendicular to each other, making the shape of a plus sign. The entire plus-maze apparatus was elevated to a height of 30 cm. Testing began by placing an animal on the central platform of the maze facing the open arm. Standard 5-min test duration was applied and the maze was wiped with 30% ethanol in between trials. Ethovision XT13 software (Noldus) was used to record the time spent on open arms and closed arms, total distance moved, and number of open arm and closed arm entries.

#### Cued threat conditioning

Mice were habituated for 15 min in the threat conditioning chambers housed inside sound attenuated cubicles (Coulbourn instruments) for two days. The habituation and training context included metal grid floor and white house light. For simple threat conditioning, mice were placed in the context for 270s and then presented twice with a 5kHz, 85 dB pure tone for 30s that co-terminated with a 2s 0.5mA footshock. The inter-trial interval (ITI) was 2 min and after the second tone-shock presentation, mice remained in the chamber for an additional 120s. Cued threat conditioning (cTC) LTM was tested 24h after training in a novel context (Context B: vanilla scented cellulose bedding, plexiglas platform and red houselight) with three presentations of paired tone (conditioned stimulus, CS). Short term memory (STM) was tested 1h after training in a different context (Context C: textured blue rubber platform, peppermint odor) with a single presentation of CS. For differential threat conditioning, mice were placed in the training context for 250s and then trained with interleaved presentations of three paired tones or CS+ (7.5 kHz pulse, 50% duty cycle) that co-terminate with 0.5mA footshock and three unpaired tones or CS− (3 kHz pure tone) in the training context with variable ITI. After the last CS− tone, mice remained in the context for an additional 140s. The following day, cued threat discrimination (cTD) LTM was tested with 3 interleaved presentations of CS+ and CS− tones with the order reversed from the training day and with variable inter-trial intervals. Freezing behavior was automatically measured by Freeze Frame software (ActiMetrics) and manually re-scored and verified by an experimeter blind to the genotype/drug.

#### Western immunoblotting

Mice were euthanized by cervical dislocation. 300 μm-thick brain slices containing amygdala [Bregma −1.22 mm to −2.06 mm] were prepared in cold (4°C) carbooxygenated (95% O_2_, 5% CO_2_) cutting solution (110 mM sucrose, 60 mM NaCl, 3 mM KCl, 1.25 mM NaH_2_PO_4_, 28 mM NaHCO_3_, 5 mM Glucose, 0.6 mM Ascorbate, 7 mM MgCl_2_ and 0.5 mM CaCl_2_) using a VT1200S vibratome (Leica). Amygdala was micro-dissected from the brain slices and sonicated in ice-cold homogenization buffer (10 mM HEPES, 150 mM NaCl, 50 mM NaF, 1 mM EDTA, 1 mM EGTA, 10 mM Na_4_P_2_O_7_, 1% Triton X-100, 0.1% SDS and 10% glycerol) that was freshly supplemented with 10 μl each of protease inhibitor (Sigma) and phosphatase inhibitor (Sigma) per ml of homogenization buffer. Protein concentrations were measured using BCA assay (GE Healthcare). Samples were prepared with 5X sample buffer (0.25 M Tris-HCl pH6.8, 10% SDS, 0.05% bromophenol blue, 50% glycerol and 25% − β mercaptoethanol) and heat denatured at 95°C for 5 min. 40 μg protein per lane was run in pre-cast 4-12% Bis-Tris gels (Invitrogen) and subjected to SDS-PAGE followed by wet gel transfer to PVDF membranes. After blocking in 5% non-fat dry milk in 0.1M PBS with 0.1% Tween-20 (PBST), membranes were probed overnight at 4°C using primary antibodies (goat anti-biotin (abcam ab53494), rabbit anti-p-eIF2α S51 (Cell Signaling #9721), rabbit anti-t-eIF2α (Cell Signaling #9722), rabbit anti-PKR (abcam ab32506, Cell Signaling #3072), rabbit ATF4 (Santa Cruz sc-390063), rabbit anti-pERK1/2 Thr202/Tyr204 (Cell Signaling #9101), rabbit anti-ERK1/2 (Cell Signaling #9102), rabbit anti-pS6 Ser(240/Ser244 (Cell Signaling #5364), mouse anti-S6 (Cell Signaling #2317), mouse anti-puromycin (Millipore MABE343), rabbit anti-Gadd34 (Proteintech #10449-1-AP), mouse anti- β tubulin (Sigma #T8328) and mouse anti- β actin (Sigma #A1978). After washing 3 times in 0.1% PBST, membranes were probed with horseradish peroxidase-conjugated secondary IgG (1:5000) (Millipore) for 1h at RT. Signals from membranes were detected with ECL chemiluminescence (Thermo Pierce) using Protein Simple instrument. Exposures were set to obtain signals at the linear range and then normalized by total protein and quantified via densitometry using ImageJ software.

#### Metabolic labeling *in vitro*

To visualize the inhibition of protein synthesis metabolic labeling experiments were conducted. 20 h post transfection with 5 μg iPKR construct (with or without NS3/4A) and 10 μL reagent cells in 6-well plate were labeled with 10 μCi TRAN^35^S label (MP Biomedicals) for 30 min following 30 min of starvation before the addition of the label into L-methionine and L-cysteine-free DMEM media (Invitrogen). Equal amounts of protein from extracts were separated by electrophoresis and transferred on a PVDF membrane. The membrane was dried in 100% methanol and analyzed for autoradiography, later it was blotted with antibodies and finally stained with Coomassie Plus stain (Pierce).

#### Bio-orthogonal non canonical amino acid tagging (BONCAT) *ex vivo*

Brain slices containing amygdala were prepared as described above and equilibrated in carboxygenated 1:1 cutting solution: ACSF (in mM 125 NaCl, 2.5 KCl, 1.25 NaH_2_PO_4_, 25 NaHCO_3_, 25 glucose, 1 MgCl_2_ and 2 CaCl_2_) for 20 min. Brain slices were further allowed to recover in ACSF at 31.2°C for 45 min, which was then supplemented with 1 mM AHA and either 1 μM ASV or vehicle for 3h. The amygdala then was micro-dissected and stored at −80°C until ready for sonication in chelator-free lysis buffer (10 mM HEPES, 150 mM NaCl, 10 mM Na4P_2_O_7_ and 1% SDS) supplemented with 20 μl each of protease and phosphatase inhibitors per ml of lysis buffer. 80 μg AHA-tagged protein lysate was subjected to cyclo addition using Protein Reaction Buffer kit with Biotin alkyne following manufacturer’s instructions (Life Technologies). Samples were prepared with 5X sample buffer and subjected to immunoblots with anti-Biotin.

#### Surface labeling of translation (SUnSET) *in vivo*

For SUnSET *in vivo* immunoblotting, Nes.iPKR mice were sequentially infused icv with 150 pg ASV (2 μl, 100 nM) infusion followed 15 min later with 25 μg puromycin. Brain slices were prepared as described above in cutting solution and the amygdala was micro-dissected and stored at −80°C until ready for homogenization in lysis buffer (10 mM HEPES, 150 mM NaCl, 50 mM NaF, 1 mM EDTA, 1 mM EGTA, 10 mM Na4P2O7, 1% Triton X-100, 0.1% SDS and 10% glycerol). 40 μg puromycylated protein lysate were subjected to Western blots with anti-puromycin. For SUnSET *in vivo* immunohistochemistry, CamK2α.iPKR mice were infused sequentially with ASV ((2 μl, 100 nM) icv followed 15 min later with puromycin (0.5 μl, 10 μg/μl) in the lateral amygdala. 1h post puromycin infusion, animals were transcardially perfused with 0.1M PBS, 0.015% digitonin followed by 4% paraformaldehyde in PBS.

#### Immunohistochemistry

Mice were deeply anesthetized with a mixture of ketamine (150 mg/kg) and xylazine (15 mg/kg), and transcardially perfused with 0.1M PBS followed by 4% paraformaldehyde in PBS. Brains were removed and postfixed in 4% PFA for 24h. 40 μm free-floating coronal brain sections containing amygdala were collected using Leica vibratome. After blocking in 5% normal goat serum in 0.1M PBS with 0.1% Triton X-100, brain sections were probed overnight with primary antibodies (chicken anti-EGFP (abcam #ab13970), rabbit anti-EGFP (Thermo Fisher #G10362), rabbit anti-p-eIF2α S51 (Cell Signaling #3398), chicken anti-mCherry (abcam #ab205402) and mouse anti-puromycin (Millipore #MABE343)). After washing three times in 0.1M PBS, brain sections were incubated with Alexa Fluor conjugated secondary antibodies (1:200) in blocking buffer for 1.5h at RT, and mounted using Prolong Gold antifade mountant with DAPI for nuclear counterstain. Imaging data were acquired using an SP8 confocal microscope (Leica) with 10x and 20X objective lenses, and analyzed with ImageJ using the Bio-Formats importer plugin. To quantify the p-eIF2α and puromycin signal intensity of each GFP immunoreactive cell, z-stacks (10 optical sections with 0.563 μm step size) for three coronal sections per mouse (n=3 mice) were collected with 20X objective with 2X zoom. All compared samples were processed using the same protocol, and images were taken with equal microscope settings. Images were analyzed using ImageJ software. To compare across groups all measures were normalized to the average intensity of the control group.

#### Statistics

Statistical analyses were performed using GraphPad Prism 8 (GraphPad software). Data are expressed as mean +/− SEM. Data from two groups were compared using two-tailed unpaired Student’s t test. Multiple group comparisons were conducted using One-way ANOVA, or Repeated Measures Two-way ANOVA, with post hoc tests as described in the appropriate figure legend. Statistical analysis was performed with an α level of 0.05. p values <0.05 were considered significant.

## Acknowlegements

We thank Maggie Marmacz for expert technical assistance, Charles Rice (The Rockefeller University) for providing the plasmid for HCV polyprotein, Martin Ryan (University of St Andrews) for the 2A plasmid, Apostolos Klinakis (Biomedical Research Foundation Academy of Athens), Ana Domingos, and Jeffrey Friedman (The Rockefeller University) for the plasmid containing the STOP sequence and the *Eef1a1* targeting plasmid, Peter Vandenabeele (Ghent University) for the PKR kinase domain plasmid and Randal Kaufman (Sanford Burnham Prebys Medical Discovery Institute) for the *Col1a1^TRE GFP.shmir4E.389^* mouse line. We thank Allen Brain Institute for providing AAV.*CAG Pr. DIO.tTA.* We thank all members of Klann lab for critical feedback and discussions. This study was supported by National Institute of Health grants NS034007 and NS047384 to E.K., Howard Hughes Medical Investigator grant to N.H., NARSAD Young Investigator grant to P.S.

## Author Contributions

P.S., P.A., N.H. conceptualized the iPKR system. P.S. and E.K. conceptual design of all *in vivo* work. P.A. designed and characterized the iPKR system *in vitro* and generated the iPKR mouse model under the supervision of N.H. P.S. carried out behavior training, and collected and analyzed *in vivo and ex vivo* data. P. M-H. carried out behavior training, F.L. carried out and analyzed slice electrophysiology, A.G. helped with mouse behavior training. J.L. provided critical advice on behavioral design. P.S. and E.K. wrote the paper. All authors read and commented on the paper.

