## Supplemental Figures for "Chemogenetic evidence that rapid neuronal *de novo* protein synthesis is required for consolidation of long-term memory"

### SUPPLEMENTAL FIGURE 1

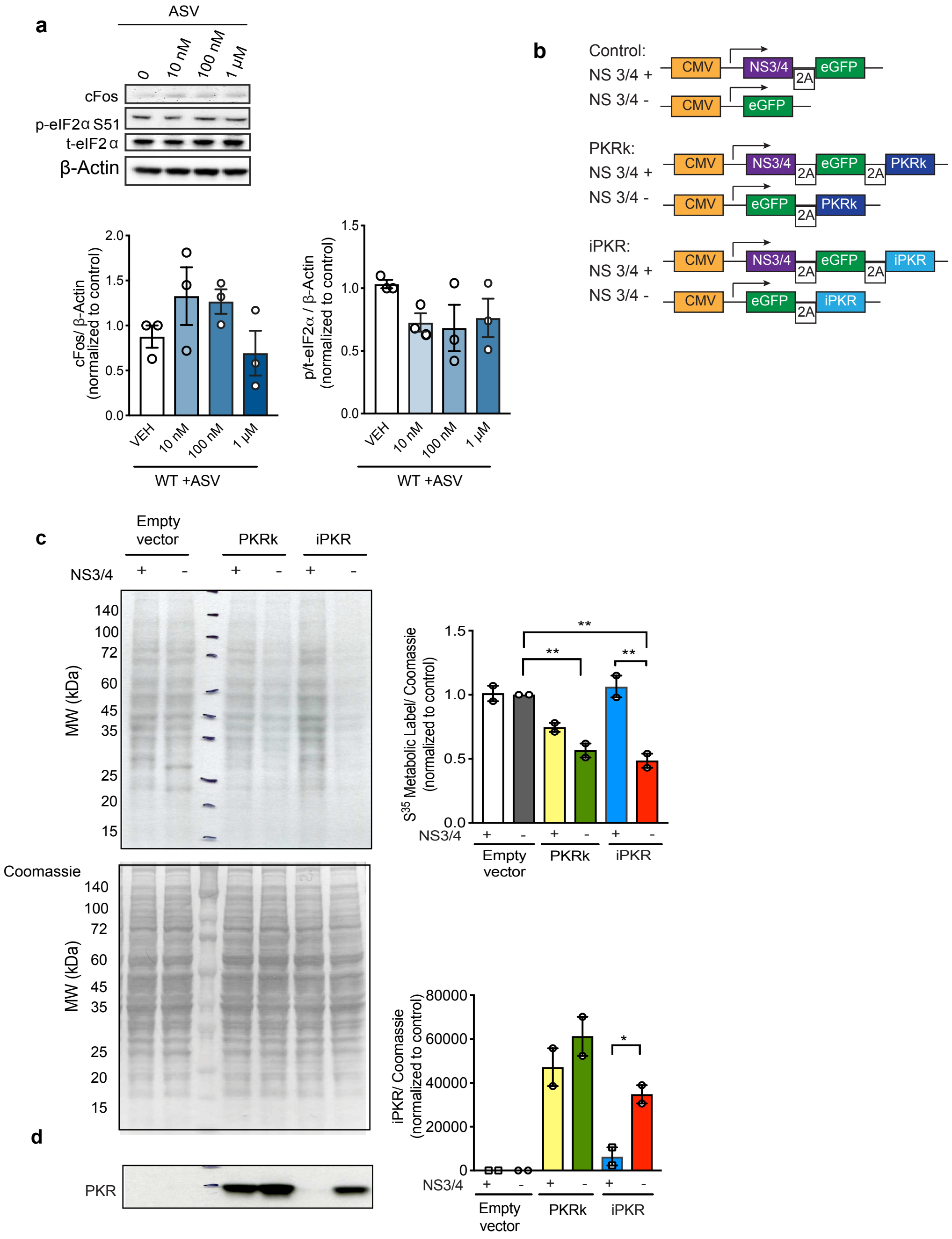


**Supplemental Figure 1.** Characterization of iPKR system. a) The toxicity of NS3/4 inhibitor ASV was determined by central i.c.v. administration of varying doses of the drug. Representative Western blots (top) shows the levels of cFos, phospho-eIF2α, total eIF2α, and control ! -Actin band in response to the administration of ASV at the doses : 0, 10 nM, 100 nM (1 µM in 2 µl saline). Bar graph with individual data points shows quantification of cFOS (left) and phospho/total eIF2α (right) normalized to ! -Actin. n=3 independent Western blots, 3 mice per group. One-way ANOVA. b) Schematic of the engineering approach for chemogenetic protein synthesis inhibitor plasmid construct consisting of NS3/4 protease, EGFP and iPKR kinase domain separated by 2A ribosome skipping sites under CMV promoter. Control plasmids harbored iPKR without NS3/4 protease or unmodified PKR kinase domain (PKRk). c) Metabolic $S^{35}$ labeling of *de novo* translation *in vitro* showed significantly decreased translation in the presence of PKRk and iPKR (**p<0.01) but the translation block was lifted by the co-expression of NS3/4 protease that degrades iPKR (**p<0.01). n = 2 biological replicate lysates per group; One way ANOVA followed by Bonferroni's post-hoc test. $F(5,6) = 19.01$, **p=0.0013. d) iPKR expression is correspondingly regulated by NS3/4 protease (**p<0.01), whereas unmodified PKRk levels were unaltered by NS3/4 protease. n = 2 per group; One way ANOVA followed by Bonferroni's post-hoc test. $F(5,6) = 22.21$, ***p=0.0008. Data are presented as +/- SEM.

### SUPPLEMENTAL FIGURE 2

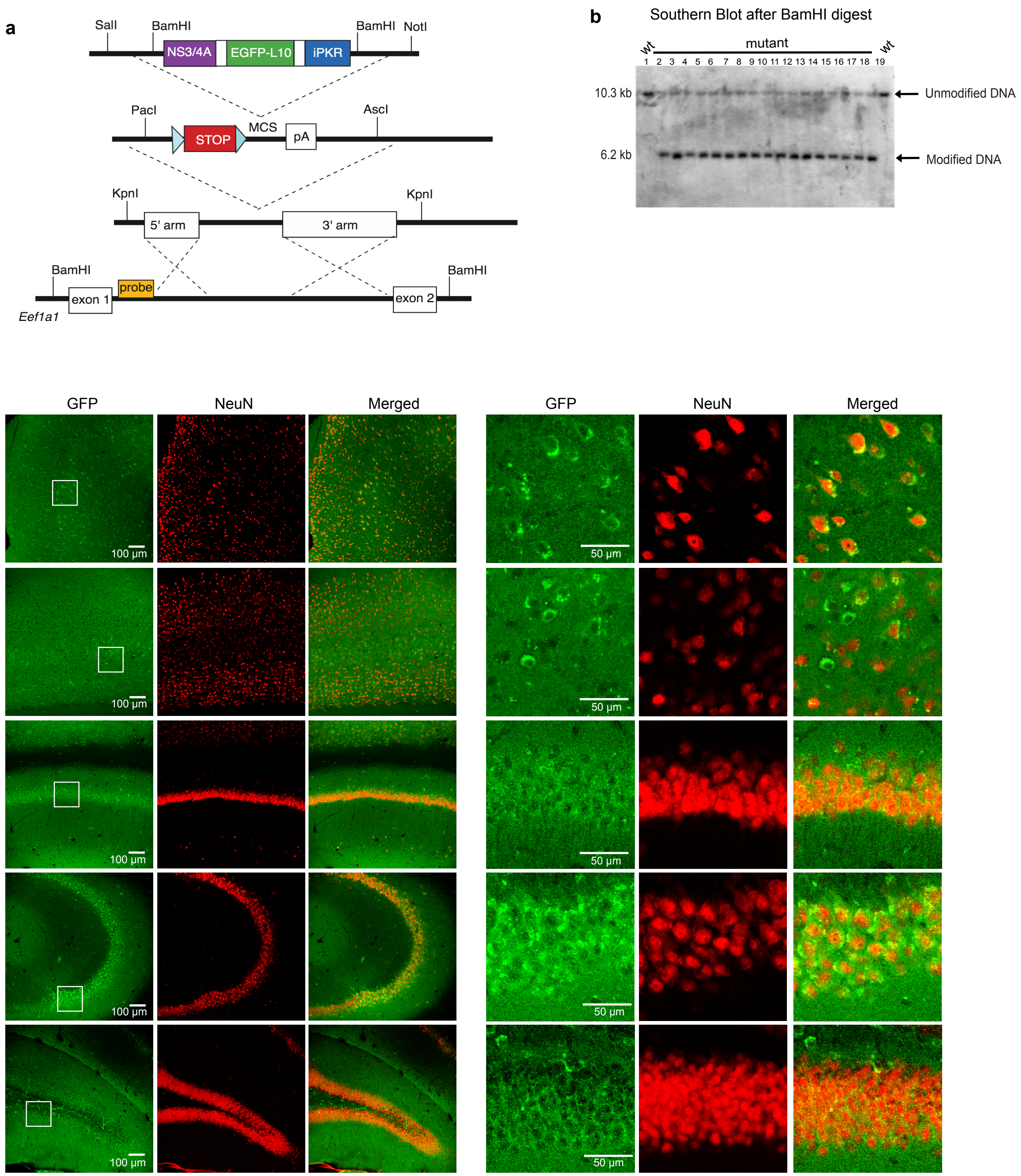


**Supplemental Figure 2.** Generation of iPKR mouse. a) Schematic showing the subcloning and targeting strategy of the multicistronic cassette containing loxP-flanked STOP cassette, NS3/4 protease, EGFP-L10, and iPKR kinase domain, which were separated by 2A ribosome skipping sites. The entire cassette was inserted between exon1 and exon 2 of *Eef1a1* genomic locus in mouse ES cells. Recognition site for the Southern blot probe is indicated. b) Southern blot after BamHI restriction enzyme-digested DNA isolated from embryonic stem cells using the probe indicated in a). Modified (6.2 kB kb) and unmodified (10.3 kB) DNA bands are indicated with arrows. In Nes iPKR brains, EGFP-L10 is expressed in the soma of neurons in the anterior cingulate cortex (c), somatosensory cortex (d), CA1 (e), CA3 (f) and dentate gyrus (g) consistent with NeuN expression. Insets show the corresponding brain areas at higher magnification.

#### SUPPLEMENTAL FIGURE 3

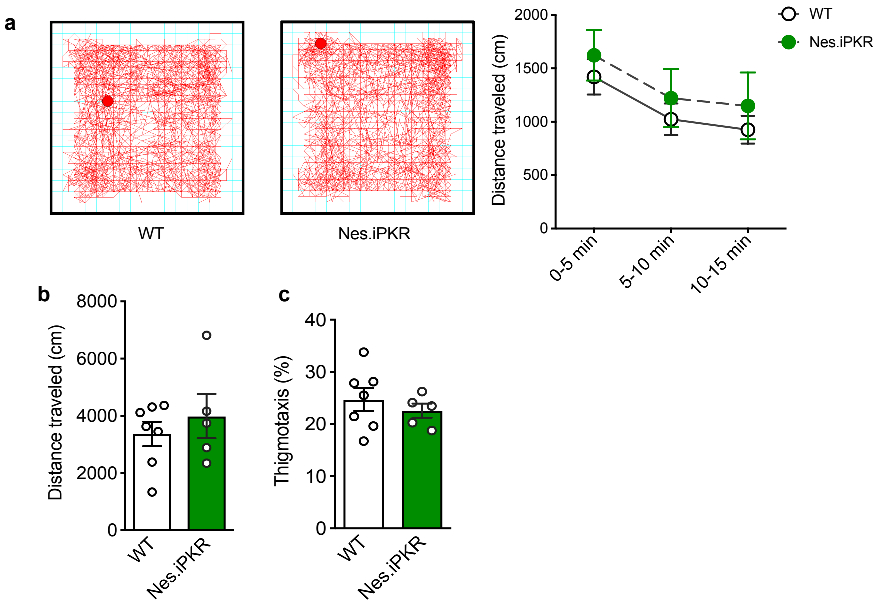


**Supplemental Figure 3.** Nes.iPKR mice display normal locomotor and anxiety related behavior. a) Nes.iPKR mice acclimated to the novel environment equivalent to the wildtype and exhibited c) normal locomotor activity in the open field test. d) Nes.iPKR animals displayed normal thigmotaxis as assessed by % distance traveled in the center compared to total distance. n = 5-7 per group; Repeated measures One-way ANOVA for a) and Unpaired t-test for c) and d).

### SUPPLEMENTAL FIGURE 4

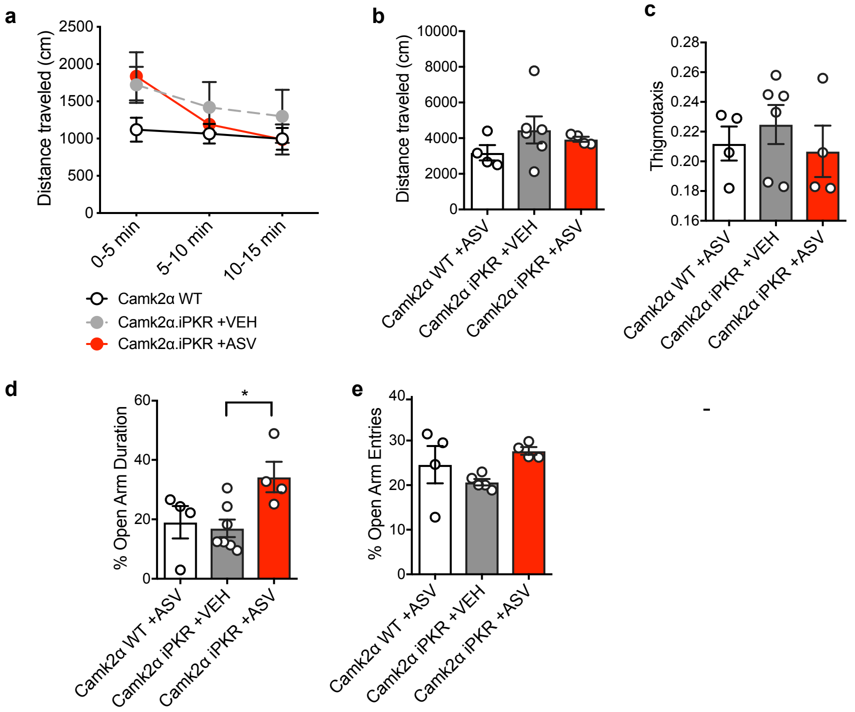


**Supplemental Figure 4.** Blocking protein synthesis in CamK2α principal neurons in LA did not affect acclimation to a novel environment (a), total locomotor activity (b), or thigmotaxis, assessed by % distance traveled in center compared to total distance (c). d) In the elevated plus maze, however, mice with protein synthesis blocked in CamK2α principal neurons exhibited reduced anxiety i.e. increased % open arm duration (*$p<0.05$) compared to vehicle (VEH)-treated CamK2α iPKR mice and CamK2α wild-type mice even though they make equivalent entries to the open arm (e). n = 4-5 per group. RM Two-way ANOVA for a), One-way ANOVA followed by Bonferroni's post-hoc test for (b), (c), (d) and (e).

### SUPPLEMENTAL FIGURE 5

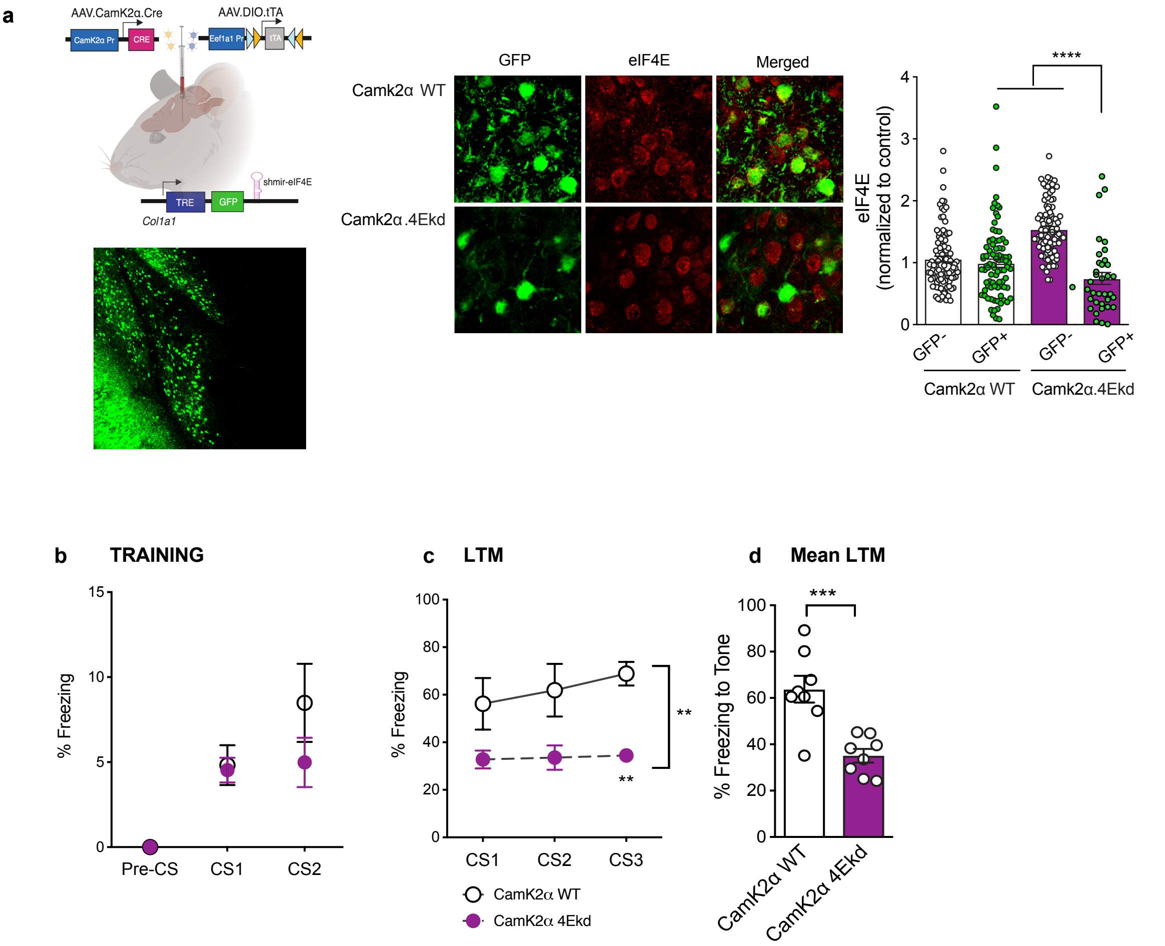


**Supplemental Figure 5.** a) Alternate strategy of blocking translation in CamK2α principal neurons in LA using cre-tet regulated synthetic micro-RNA targeted against eIF4E. Col1a1.TRE.GFP.shmir4E mice were bilaterally injected in the lateral amygdala with AAV1.CamK2α.Cre and AAV9.DIO.tTA, and placed off dox diet for 10 days before training. b) eIF4E protein level was significantly decreased in GFP+ neurons that express shmir4E. Two-way ANOVA with Bonferroni's post-hoc test. Genotype X GFP interaction: $F(1,311) = 32.29$, ****$p<0.0001$; GFP: $F(1,311) = 45.32$, ****$p<0.0001$. c) CamK2α 4Ekd mice learned the association between CS and US during training. RM Two-way ANOVA with Bonferroni's post-hoc test. CS: $F(2,14) = 17.54$, ***$p=0.0002$. d) Cued LTM was severely impaired across all three CS presentations. n=8 per group; RM Two-way ANOVA with Bonferroni's post-hoc test. $F(10,20) = 15.65$, **$p=0.0027$. d) Mean cTC LTM was significantly impaired in CamK2α 4Ekd mice compared to wildtype (***$p<0.001$). n = 8 per group; Unpaired t-test. Data are presented as +/- SEM.

### SUPPLEMENTAL FIGURE 6

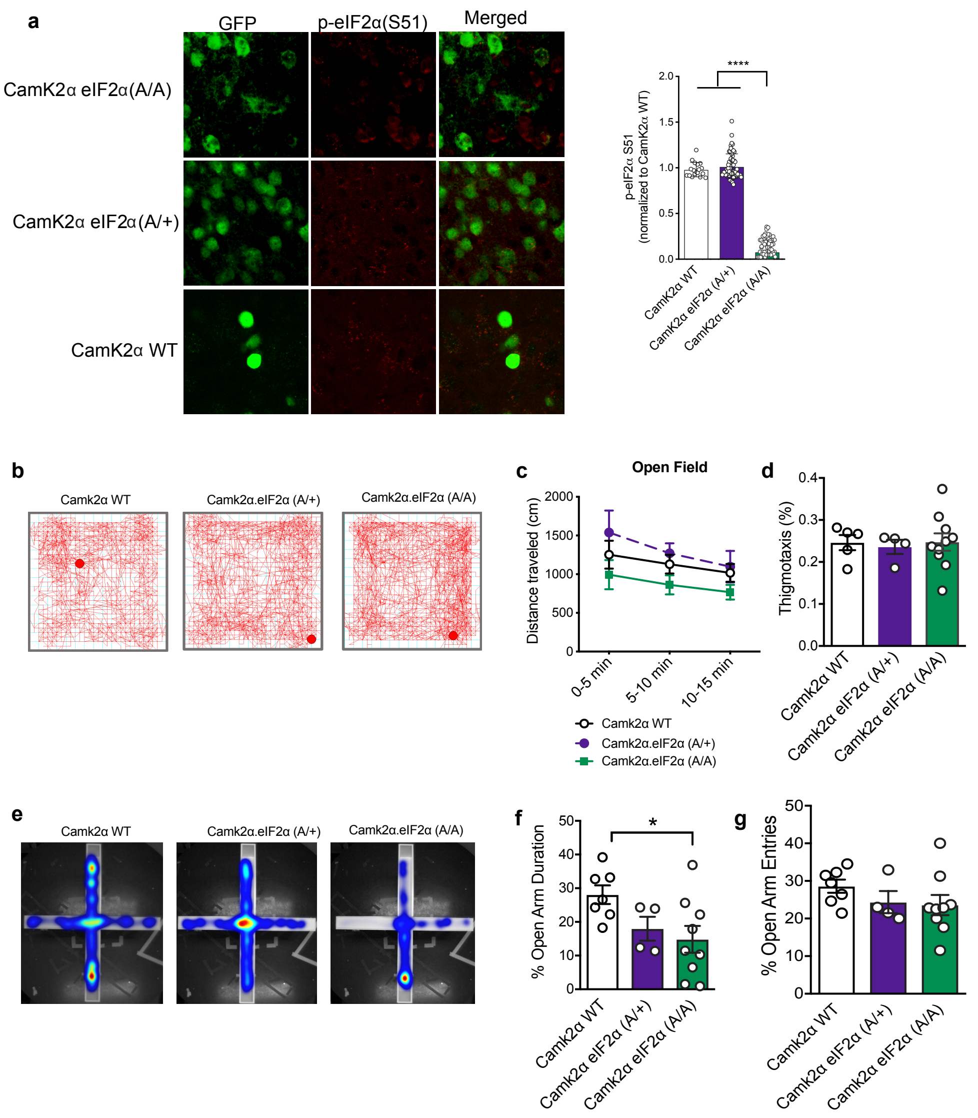


**Supplemental Figure 6.** a) eIF2α phosphorylation at S51 was significantly reduced in GFP+ neurons in CamK2α.eIF2α (A/A) mice compared to GFP- neurons, as well as GFP+ neurons in CamK2α.WT mice (****p<0.001). (n = 60-74 per group, 3 animals); One-way ANOVA with Bonferroni's post-hoc test. $F(2, 154) = 1055$, ****$p<0.0001$. b) Representative motion traces from the open field test for CamK2α.WT, CamK2α.eIF2α(A/+) and CamK2α eIF2α (A/A) mice. c) In the open field test, CamK2α.eIF2α (A/A) mice acclimated to the novel environment and had comparable spontaneous locomotion compared to the CamK2α.WT mice and CamK2α. eIF2α (A/+) mice. RM One-way ANOVA. d) Bar graphs representing thigmotaxis, i.e. %time spent in center compared to total distance traversed in the open field arena for the three groups. One-way ANOVA. f) In the elevated plus maze, CamK2α.eIF2α (A/A) mice spent a significantly higher duration in the open arm compared to CamK2α WT mice (g) (*p<0.05) indicating anxiety like behavior, even though they made equivalent entries to the open arm (h). One-way ANOVA with Bonferroni's post-hoc test. $F(2,17) = 3.775$, *$p=0.0440$. Data are presented as +/- SEM

### SUPPLEMENTAL FIGURE 7

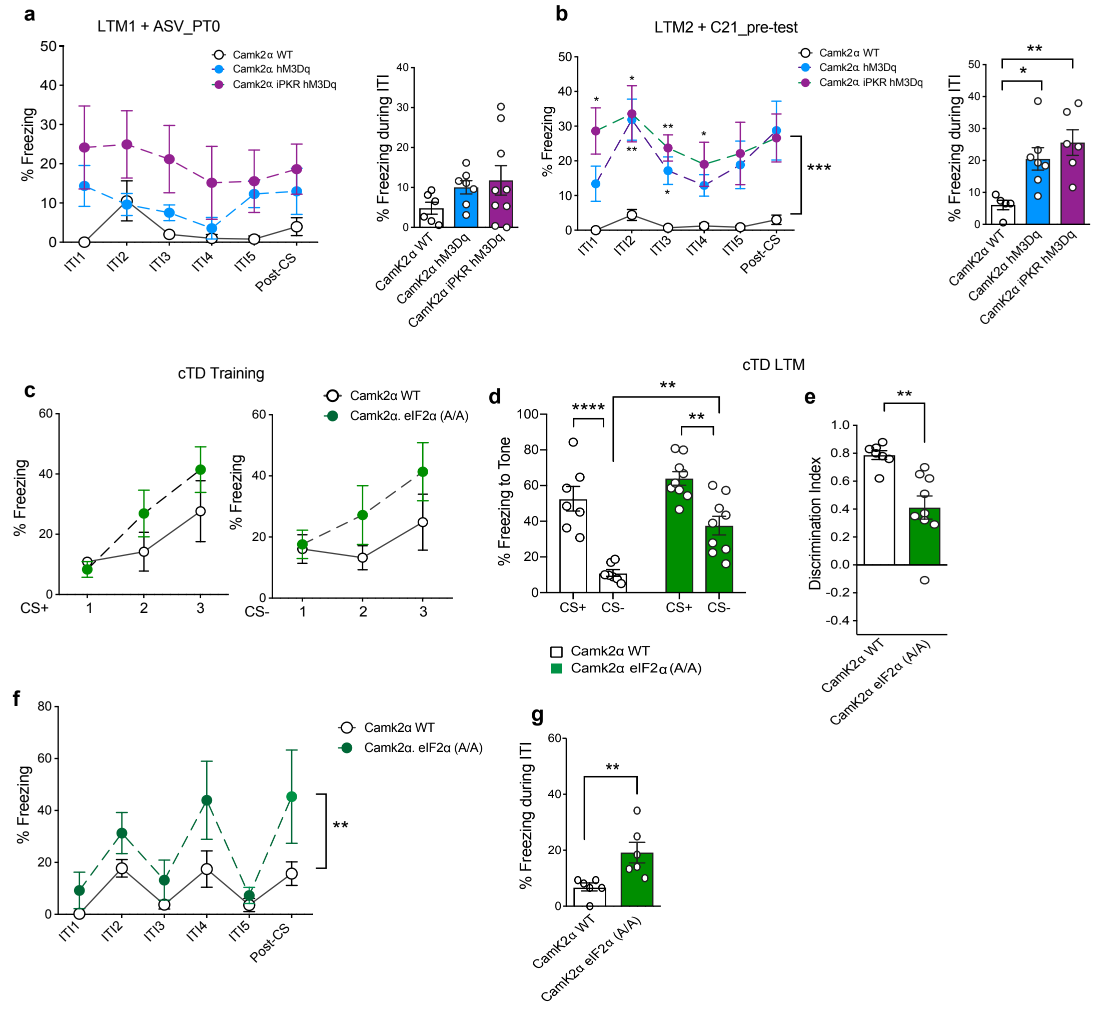


**Supplemental Figure 7.** a) All groups of mice - CamK2α. iPKR hM3Dq, CamK2α.hM3Dq and CamK2α WT, exhibited low freezing during ITI in LTM1. XY plots showing %freezing during individual ITIs and Post-CS (left; RM Two-way ANOVA) and bar graphs showing mean %freezing during ITI (right; One-way ANOVA). n = 6-9 per group. b) During LTM2, administration of DREADD agonist C21 caused an increase in freezing during ITI for both CamK2α.hM3Dq and CamK2α. iPKR hM3Dq groups compared to CamK2α WT mice. XY plots showing %freezing during individual ITIs and Post-CS (left). n = 5-7 per group. RM Two-way ANOVA genotype: $F(2,15)=12.63$, ***$p=0.0006$). Bar graphs showing mean %freezing during ITI (right). One-way ANOVA with Bonferroni's post-hoc test. *$p<0.05$ and **$p<0.01$. c) CamK2α. eIF2α (A/A) mice displayed comparable learning in the diferential threat conditioning training for both CS+ (right) and CS- (left). d) However, in the LTM test, they displayed significant increase in CS- response compared to CamK2α WT mice (**$p<0.01$). Two-way ANOVA with Bonferroni's post-hoc test. CS: $F(1,28) = 49.18$, ****$p<0.0001$; Genotype: $F(1,28) = 15.26$, ***$p=0.0005$. e) The cTD discrimination index was significantly lower for CamK2α. eIF2α (A/A) mice (**$p<0.01$) relative to controls. n=7-10 per group; Unpaired t-test. f) Besides stimulus generalization, CamK2α. eIF2α (A/A) mice also displayed cognitive inflexibility and could not stop freezing after the tone offset, and thus had significantly higher freezing rate during the ITIs. RM Two-way ANOVA with Bonferroni's post-hoc test. Genotype: $F(1,10) = 16.70$, **$p=0.0022$. g) Mean freezing response during ITI is significantly increased in CamK2α. eIF2α (A/A) mice (**$p=0.0097$) . n=7-10 per group; Unpaired t-test. Data are presented as mean +/- SEM.
